# A Spatiotemporal Molecular Switch Governs Plant Asymmetric Cell Division

**DOI:** 10.1101/2020.09.05.284380

**Authors:** Xiaoyu Guo, Chan Ho Park, Zhi-Yong Wang, Bryce E. Nickels, Juan Dong

## Abstract

Asymmetric cell division (ACD) often requires protein polarization in the mother cell to produce daughter cells with distinct identities (“cell-fate asymmetry”). Here, we define a previously undocumented mechanism for establishing cell-fate asymmetry in *Arabidopsis* stomatal stem cells. In particular, we show that polarization of BSL protein phosphatases promotes stomatal ACD by establishing a “kinase-based signaling asymmetry” in the two daughter cells. BSL polarization in the stomatal ACD mother cell is triggered upon commitment to cell division. Polarized BSL is inherited by the differentiating daughter cell where it suppresses cell division and promotes cell-fate determination. Plants lacking BSL exhibit stomatal over-proliferation, demonstrating BSL plays an essential role in stomatal development. Our findings establish that BSL polarization provides a spatiotemporal molecular switch that enables cell-fate asymmetry in stomatal ACD daughter cells. We propose BSL polarization is triggered by an ACD “checkpoint” in the mother cell that monitors establishment of division-plane asymmetry.

## Introduction

During the development of multicellular organisms, stem cells undergo self-renewing asymmetric cell division (ACD) to generate diverse cell types (Abrash and Bergmann, 2009; Knoblich, 2008). In ACD, the division of a progenitor cell (“mother cell”) yields one daughter cell identical to the progenitor and one “differentiating daughter cell” that adopts a distinct identity. Thus, in contrast to symmetric cell division, which yields two daughter cells identical to the progenitor, ACD requires cellular processes that establish “cell-fate asymmetry” (Pierre-Jerome et al., 2018; Ruijtenberg and van den Heuvel, 2016). The asymmetric distribution, or “polarization,” of factors in the progenitor cell provides a mechanism by which cell-fate determinants can be asymmetrically distributed between daughter cells. Protein polarization enables cell-fate determinants to be sequestered in a location within the progenitor cell that is partitioned to one of the two daughter cells after division. Accordingly, establishment of cell-fate asymmetry often requires protein polarization in ACD progenitor cells (Goldstein and Macara, 2007; Sunchu and Cabernard, 2020).

Stomatal development in *Arabidopsis* provides a model system for understanding how protein polarization in an ACD progenitor cell results in cell-fate asymmetry in daughter cells. Stomatal development begins with the emergence of a stomatal-lineage stem cell, “meristemoid mother cell (MMC),” from a subset of protodermal cells (PrC) located in the epidermis of the developing leaf (Figure 1A, left). The MMC undergoes ACD, yielding two asymmetric daughter cells: a smaller meristemoid (M) and a larger stomatal-lineage ground cell (SLGC) (Facette and Smith, 2012). The meristemoid (M) undergoes subsequent rounds of cell division prior to differentiating into guard cells (GCs), two of which comprise a stomatal pore that facilitates exchange of gas and water vapor between the plant and the environment (Figure 1A, right). The SLGC, in contrast to the M, has a limited ability to divide and typically differentiates into a non-stomatal pavement cell (PC), which provides protection and structural support for other cell types in the leaf. Stomatal divisions can be responsive to external cues, thus enabling plants to optimize the number and distribution of stomatal pores in response to environmental or developmental changes (Endo and Torii, 2019; Pillitteri and Torii, 2012; Zoulias et al., 2018).

**Figure 1.**
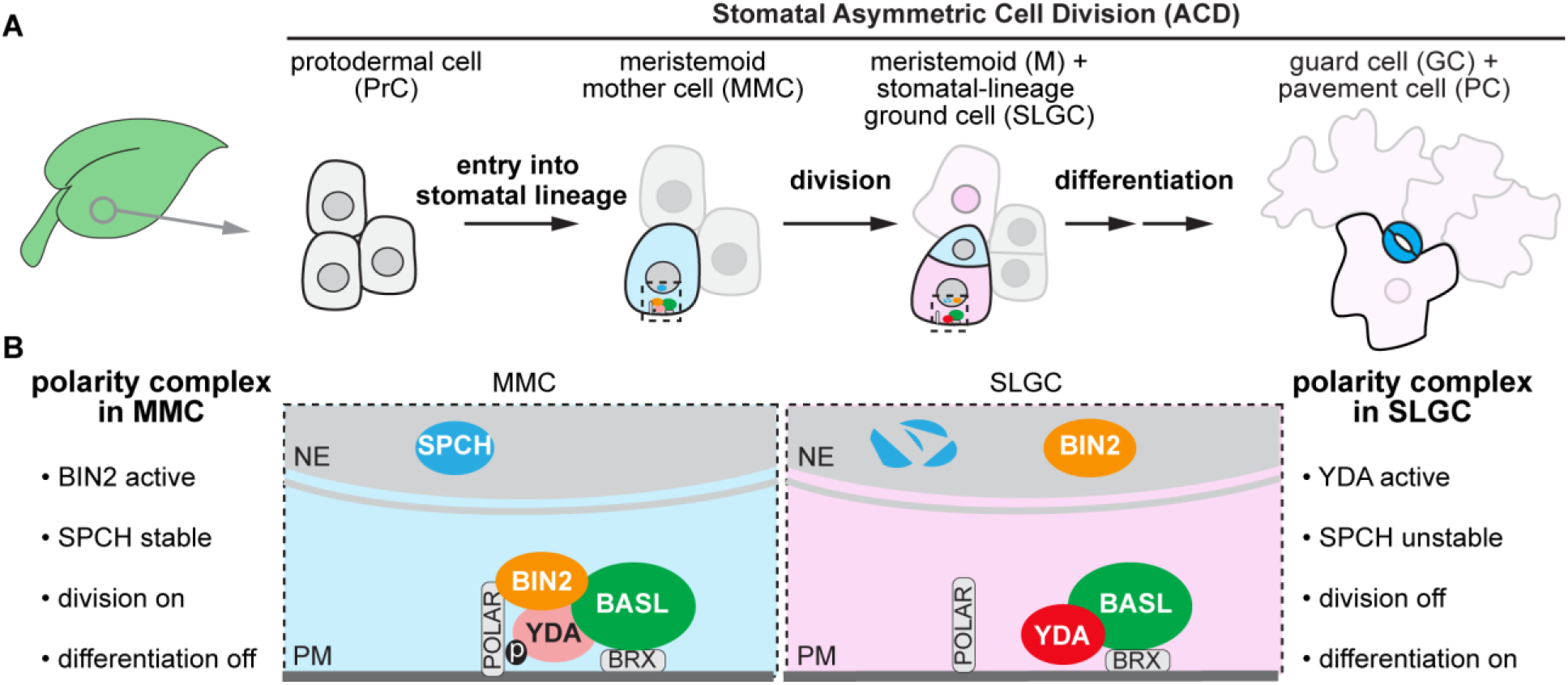
Compositional and Functional Changes of the Polarity Complex before and after a Stomatal ACD. (A) Stomatal development in a young leaf of *Arabidopsis.* Light blue, stomatal fate; dark blue, stomatal guard cells; pink, non-stomatal fate; light pink, pavement cell. Fading cells, PrCs not converted to MMC become non-stomatal pavement cells. Dotted rectangle, regions enlarged in (B) containing protein components of the polarity complex. (B) Enlarged view of polarity protein complexes required for the progenitor cell (MMC, light blue) and the daughter cell (SLGC, pink), respectively, in stomatal ACD. See also Figure S1.

Emergence of an ACD progenitor cell (MMC) from a PrC depends on the activity of the transcription factor SPEECHLESS (SPCH) (MacAlister et al., 2007), which controls the expression of genes that function in stomatal-lineage cell division and stomatal formation (Lau et al., 2014). SPCH regulates stomatal ACD, at least in part, by activating expression of the scaffold protein BASL (BREAKING OF ASYMMETRY IN THE STOMATAL LINEAGE) (Lau et al., 2014). In the MMC, BASL polarizes at the cell membrane (Figure 1A, dashed box), establishing the cellular asymmetry that ultimately leads to division-plane asymmetry (Dong et al., 2009). Polarized BASL also provides a scaffold for assembly of a “polarity complex” that is maintained during the MMC division and inherited by the SLGC daughter cell (Figure 1B).

During the progression of stomatal ACD, changes in the composition of the polarity complex facilitate cell-fate asymmetry by modulating the activity of SPCH (Figure 1B) (Houbaert et al., 2018; Zhang et al., 2015). In the MMC, the polarity complex contains the MAPKK Kinase YODA (YDA) and the GSK3-like BIN2 kinases, which function as negative and positive regulators of SPCH, respectively (Bergmann et al., 2004; Gudesblat et al., 2012; Kim et al., 2012; Lampard et al., 2008) (Figure 1B, left). YDA inhibits SPCH by stimulating a MAPK signaling cascade that results in the degradation of SPCH in the nucleus (Lampard et al., 2008). In contrast, BIN2’s association with the polarity complex inhibits YDA. Thus, BIN2 functions as a *de facto* activator of SPCH by inhibiting its degradation (Houbaert et al., 2018; Kim et al., 2012). Therefore, in the MMC, the association of BIN2 with the polarity complex enables SPCH to promote cell division (Vatén et al., 2018). In the differentiated daughter cell (SLGC), BIN2 is repartitioned to the nucleus, which relieves BIN2-dependent inhibition of YDA, resulting in the degradation of SPCH (Figure 1B, right) (Houbaert et al., 2018). Therefore, in the SLGC, the repartitioning of BIN2 to the nucleus suppresses cell division. Accordingly, differences between the composition of the polarity complex in the MMC *vs*. in the SLGC alter the cell-division potential and fate specification of these cell types (Guo and Dong, 2019). Nevertheless, the mechanistic basis for the changes in the composition of the polarity complex that are essential for cell-fate asymmetry in stomatal ACD has not been defined.

## Results

### BSL Protein Phosphatases Interact with BASL

We hypothesized that the repartitioning of BIN2 to the nucleus is driven by one or more factors that associate with the polarity complex. Therefore, we used an unbiased biochemical approach to isolate proteins in *Arabidopsis* cell extracts that associate with the polarity-complex by interacting with BASL (Figure 1). We expressed a GFP-BASL fusion or GFP alone in *Arabidopsis* and used mass spectrometry (MS) to identify candidate BASL-interacting proteins as those recovered by co-immunoprecipitation (co-IP) with GFP-BASL but not GFP (Figure 2A). We identified several proteins previously shown to associate with the polarity complex, including the BRX proteins (Rowe et al., 2019) (Figure 2A), thus validating the approach. We also identified several proteins not previously shown to associate with the polarity complex, including several members of the BSL family of Ser/Thr-protein phosphatases (BSL1, BSL2, BSL3; Figure 2A). BSL proteins were of particular interest because they have previously been shown to modulate the activity of BIN2 in plant responses to the phytohormone Brassinosteroids (BR) (Kim et al., 2009) (Figures S1A and S1B). To confirm that BSL1, BSL2, and BSL3 interact with BASL we performed pairwise yeast two-hybrid assays and *in vitro* co-IP assays using purified fusion proteins produced by *N. benthamiana* leaf cells. The results establish that BSL1, BSL2, and BSL3 directly interact with BASL, whereas we did not detect interactions between BASL and the fourth BSL protein, BSU1 (Figures 2B and S2A). Taken together, the results identify BSL1, BSL2, and BSL3 as previously unknown BASL-interacting proteins.

**Figure 2.**
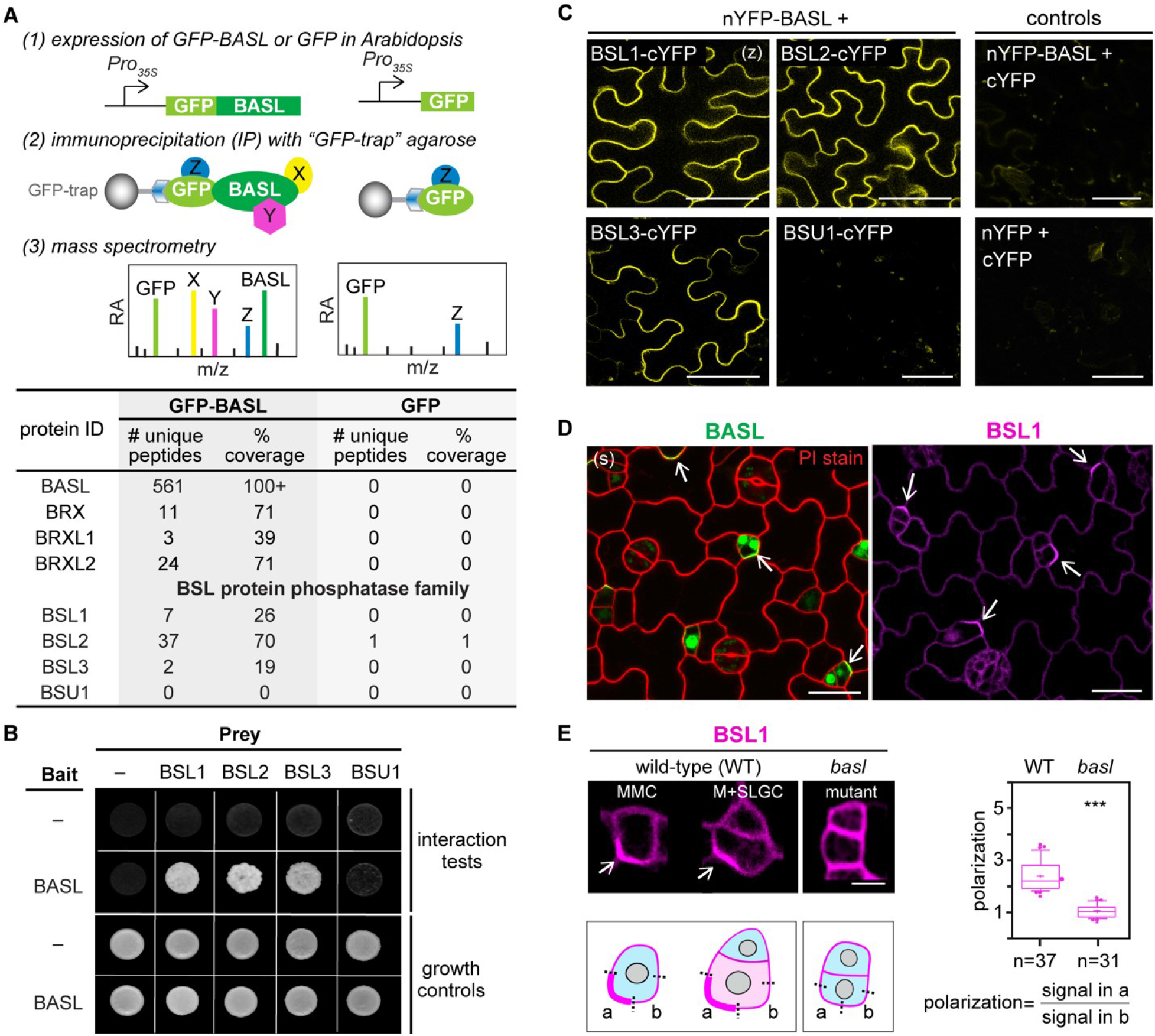
Identification of BSL Proteins as Putative Components of the BASL Polarity Complex in *Arabidopsis*. (A) Identification of BASL-interacting proteins in *Arabidopsis* using co-immunoprecipitation (IP) coupled with mass spectrometry. GFP along was used as control. Top, experimental procedure. Bottom, results. (B) BSL proteins directly interact with BASL. Results of yeast two-hybrid assays. Bait, “-” indicates Gal4 DNA-binding domain (BD) while BASL indicates BD-BASL fusion protein; Prey, “-” indicates Gal4 activation domain (AD) while BSL1, BSL2, BSL3, and BSU indicate BSL protein-AD fusions; interaction tests, assays performed using synthetic dropout medium; growth controls, assays performed using rich media. (C) BSL proteins interact with BASL at the cell membrane. Results of bimolecular fluorescence complementation (BiFC) assays in *N. benthamiana* leaf epidermal cells. nYFP, N-terminal YFP; cYFP, C-terminal YFP. YFP signals indicate protein-protein interactions. Scale, 50 μm. (z), z-staked confocal images. (D) BSL1 polarizes in stomatal lineage cells. Images show cells expressing GFP-BASL (green) or BSL1-YFP (magenta), each driven by their endogenous promoter. Arrows indicate protein polarization. Red, PI staining. Scale, 20 μm. (E) BSL1 polarization requires BASL. Left, localization of BSL1-YFP in wild-type or *basl-2*. Arrows indicate protein polarization. Scale, 2 μm. Polarization is calculated as the ratio of the fluorescence in segment a *vs.* b. Right, quantitation of BSL1 polarization. Box plot shows first and third quartiles, median (line) and mean (cross). n, number of cells. ***P < 0.0001. See also Figure S2.

To determine where BSL1, BSL2, and BSL3 interact with BASL in plant cells, we performed bimolecular fluorescence complementation (BiFC) assays in *N. benthamiana* leaf epidermal cell (Figures 2C, S2B and S2C). The BiFC assay relies on the ability of non-fluorescent fragments of YFP (nYFP and cYFP) to complement one other when brought within close proximity. Thus, we monitored the fluorescent signal observed in cells in which nYFP-tagged BASL was co-expressed with cYFP-tagged BSL. Strikingly, we observe a strong fluorescent signal associated with the cell membrane in plants co-expressing nYFP-BASL and cYFP-tagged BSL1, BSL2, or BSL3 (Figures 2C and S2C). In contrast, no fluorescence was detected in plants co-expressing nYFP-BASL and BSU1-cYFP (Figures 2C and S2C). Analysis of the subcellular localization of full-length YFP-tagged BASL, BSL1, BSL2, or BSL3 indicates that, when expressed in the absence of an interacting partner, these proteins are distributed in the cytoplasm and nucleus (Figure S2B). Thus, results of BiFC complementation assays indicate that interactions between BASL and BSL1, BSL2, or BSL3 proteins stabilize the association of these factors with the cell membrane in plants.

### BSL Proteins Co-Localize with Polarized BASL in Stomatal-Lineage Cells

To establish whether BSL proteins interact with BASL at the cell membrane in stomatal-lineage cells, we monitored the subcellular localization of BSL-YFP fusion proteins in *Arabidopsis* leaf epidermal cells. Across all cell types, BSL1 is predominantly located in the cytoplasm close to the cell periphery while BSL2 and BSL3 are located in both the nucleus and cytoplasm (Figure S2D). However, in the subset of epidermal cells that comprise the stomatal lineage, BSL1 is polarized at the cell membrane (Figure 2D), exhibiting a localization pattern that bears a strong resemblance to the localization pattern observed for polarized BASL (Figure 2D, left). Differing from BASL, BSL1 does not localize to the nucleus where BASL is initially stored (Zhang et al., 2016). To directly determine whether BSL1 co-localizes with polarized BASL in stomatal-lineage cells, we monitored protein localization in plants co-expressing GFP-BASL and BSL1-RFP (Figure S2E). The results show, definitively, that BSL1 and BASL co-localize at the cell membrane in stomatal-lineage cells (Figures S2E and S2F). Furthermore, analysis of protein localization in plants co-expressing GFP-BASL and BSL2-RFP or BSL3-RFP show BSL2 and BSL3 also colocalize with polarized BASL in stomatal-lineage cells (Figure S2F). We conclude that BSL proteins co-localize with polarized BASL at the cell membrane in *Arabidopsis* stomatal-lineage cells.

### BSL Proteins Are Components of the “Polarity Complex” the Forms During Stomatal ACD

To determine whether BSL polarization in stomatal-lineage cells requires BASL, we monitored the localization of native promoter driven BSL1-YFP in wild-type plants or plants containing a loss-of-function *basl-2* allele. For each cell containing detectable levels of BSL1-YFP, we obtained a measure of BSL1 polarization by calculating the ratio of BSL1-YFP fluorescence at the polarity crescent (Figure 2E, box “a”) relative to the fluorescence observed in a segment of the cell membrane adjacent to the polarity crescent and with the same length (Figure 2E, box “b”). Thus, a polarization value of ~1 indicates BSL1 is not polarized while a polarization value >1 indicates BSL1 polarization. The results in Figure 2E show BSL1 polarization occurs in stomatal-lineage cells from wild-type plants (mean = 2.3, SD = 0.6, n = 37) but does not occur in cells from *basl-2* plants (mean = 1.0; SD = 0.2; n = 31). Thus, BSL1 polarization in stomatal-lineage cells requires BASL.

Taken together, the results in Figure 2 establish that BSL proteins polarize in stomatal-lineage cells, at least in part, by directly interacting with the BASL scaffold protein. Thus, BSL proteins exhibit the defining hallmark of polarity-complex associated factors. Accordingly, we conclude that BSL proteins associate with the polarity complex required in stomatal-lineage cells.

### Association of BSL1 with the Polarity Complex Occurs in MMCs Committed to Cell Division

Unexpectedly, analysis of MMCs in plants co-expressing GFP-BASL and BSL1-RFP we observed two populations of cells; one in which only GFP-BASL was polarized and another in which both GFP-BASL and BSL1-RFP were polarized (Figures 3A and S2G). Thus, BASL polarization is necessary, but is not sufficient, for BSL1 polarization. Comparison of the size and shape of the cells within each population revealed cells in which only BASL was polarized were smaller and symmetric, whereas cells in which both BASL and BSL1 were polarized were larger and asymmetric. The differences in the size and shape of cells comprising each population raises the possibility that BSL1 polarization occurs at a later time in development than BASL polarization. Therefore, to examine the timing of BASL polarization relative to BSL1 polarization, we took advantage of our ability to identify the subset of MMCs committed to cell division using the microtubule marker mCherry-TUA5 (Gutierrez et al., 2009), which enables visualization of the “pre-prophase band (PPB)” that forms at the onset of mitosis (Livanos and Muller, 2019) (Figures S3A and S3B). We used the mCherry-TUA5 marker to monitor the progression of the cell-cycle in MMCs containing polarized GFP-BASL or polarized BSL1-YFP (Figures 3B-3D). The results indicate that commitment to cell division had occurred within half of the MMCs containing polarized BASL (~52%, n = 102) but nearly all of the MMCs containing polarized BSL1 (~98%, n = 190). The results establish that BASL polarization, but not BSL1 polarization, can occur prior to the commitment of the MMC to cell division. Thus, BASL polarization occurs at an earlier developmental timepoint than BSL polarization.

**Figure 3.**
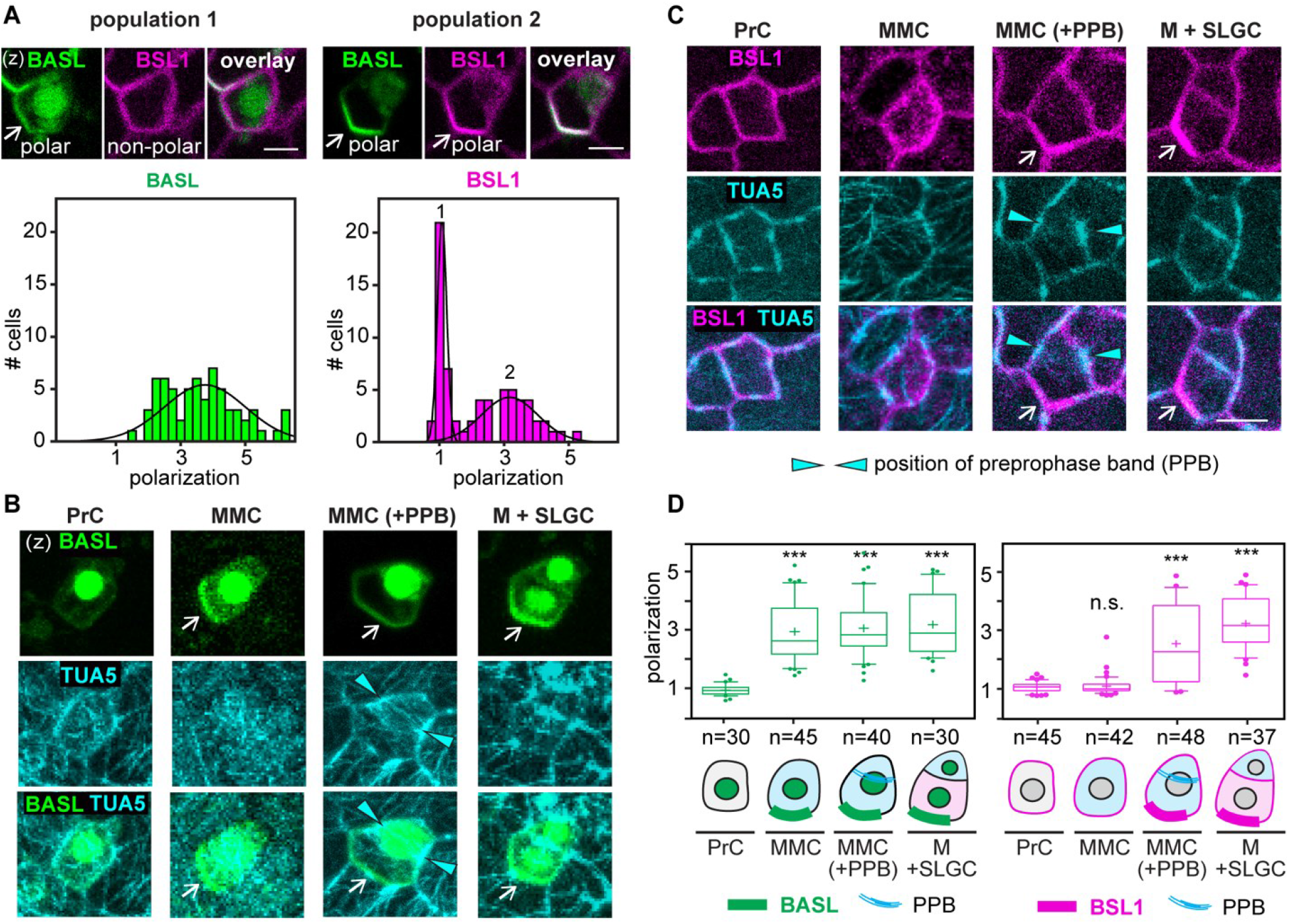
Association of BSL1 with the Polarity Complex Occurs in MMCs Committed to Cell Division. (A) Differential polarization of BASL and BSL1 in MMCs. Top, representative images of cells in indicated populations of MMCs containing BSL1-RFP (magenta) and GFP-BASL (green). Scale, 2 μm. Bottom, histograms of polarization values for BASL (left) and BSL1 (right). The two curves correspond to the two populations demonstrated above. n = 68 cell co-expressing GFP-BASL and BSL1-RFP. (B-C) BSL1 polarization coincides with formation of the pre-prophase band (PPB). Images show co-expression of GFP-BASL (B) or BSL1-YFP (C) with the microtubule marker mCherry-TUA5 (cyan) that allows visualization of the formation of the preprophase band (PPB) during mitosis. Arrow heads, PPB position. Arrows, protein polarization. Scale, 5 μm. (D) Quantification of protein polarization of BASL (left) and BSL1 (right) in successive cell types (PrC, MMC, MMC with PPB, and SLGC) during stomatal development. One-way ANOVA with Tukey’s post hoc test was performed to compare the values for designated cell type with the values for PrC. n, # of cells counted. ***P < 0.0001; n.s., not significant. See also Figure S3.

Next, to examine the timing of BSL1 polarization relative to PPB formation we selected MMCs containing the PPB and determined the percentage of these in which BSL1 was polarized. The results show that ~98% of MMCs containing the PPB also contain polarized BSL1 (n = 146). Thus, we observe a nearly one-to-one correspondence between the presence of the PPB and the presence of polarized BSL1 in MMCs. The results suggest BSL1 polarization occurs at the same time in development as PPB formation. Accordingly, we conclude that the association of BSL1 with the polarity complex occurs upon commitment of MMCs to cell division.

### Association of BSL1 with the Polarity Complex Modulates BIN2 Partitioning to the Nucleus by Disrupting BIN2-BASL Interactions

As mentioned above, repartitioning of the GSK3-like BIN2 kinases to the nucleus enables YDA-dependent suppression of SPCH and inhibition of cell division in the SLGC (Figure 1B). To determine whether BSL proteins function in the repartitioning of BIN2, we measured the ability of BSL proteins to modulate BIN2 repartitioning in stomatal-lineage cells. To do this, we analyzed the distribution of BIN2-YFP in wild-type plants, plants lacking all four BSL proteins (*bsl*-quad), or plants in which BSL1 is overexpressed (BSL1 ++) (Figures 4A and S4A-S4D). We observe a substantial decrease in the proportion of BIN2 in the nucleus in plants lacking BSL while, in contrast, we observe a substantial increase in the proportion of BIN2 in the nucleus in plants in which BSL1 is overexpressed (Figures 4A, and S4A-S4D). In addition, the changes in BIN2 localization was not recapitulated by overexpression of a catalytically inactive BSL1 variant (Mora-Garcia et al., 2004) (BSL1^D584N^) (Figure S4A), indicating that the BSL1 phosphatase activity is required to promote BIN2 repartitioning. The results establish that BSL proteins modulate the partitioning of BIN2 between the nucleus and cell membrane in stomatal-lineage cells.

**Figure 4.**
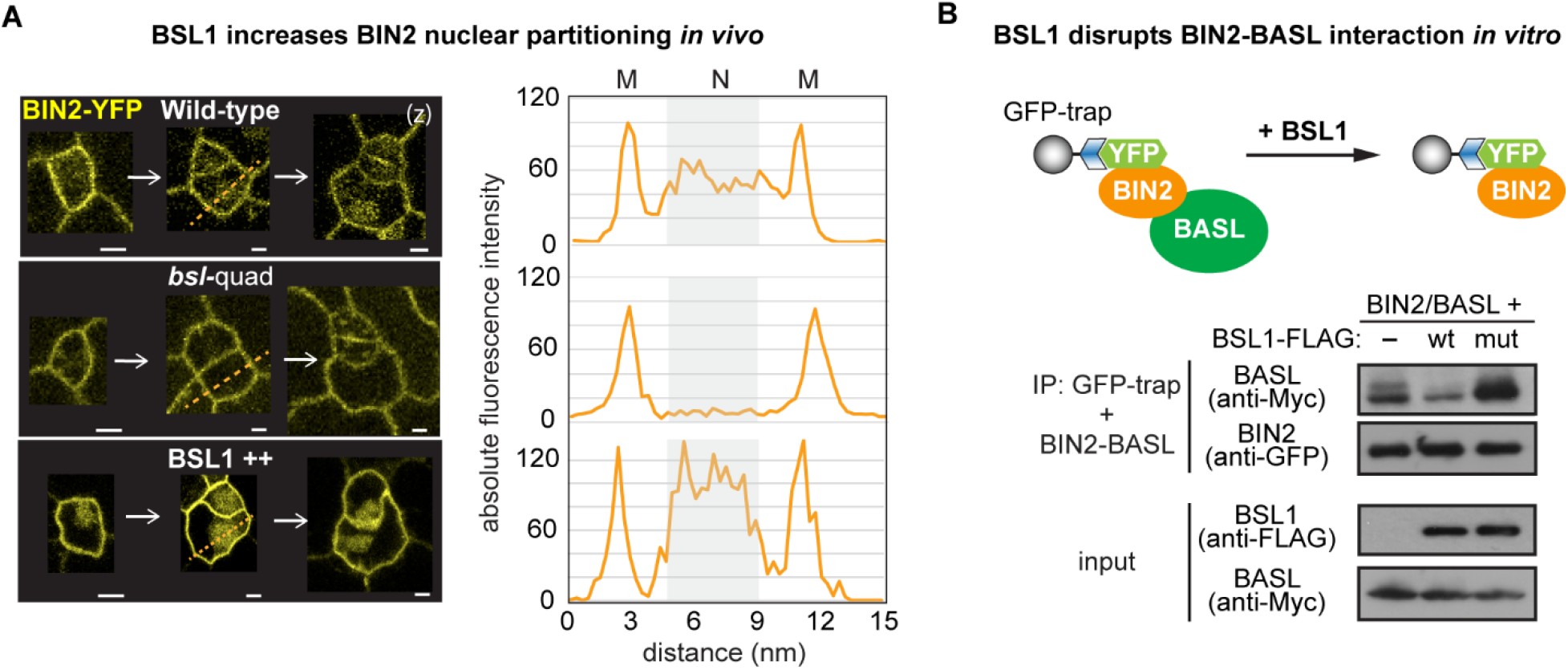
Association of BSL1 with the Polarity Complex Modulates BIN2 Partitioning to the Nucleus. (A) BSL1 facilitates the redistribution of BIN2 from the cell membrane to the nucleus. Left, BIN2-YFP localization (yellow) in the indicated genetic backgrounds. Dotted line indicates regions used for florescence intensity profiling shown on the right*. bsl*-quad, quadruple loss-of-function mutant; BSL1 ++, *BSL1* overexpression. M, cell membrane; N, nucleus. Scale, 5 μm. (B) BSL1 interferes with BIN2’s interaction with BASL. Top, schematic of co-IP assay used to monitor the ability of BSL1-FLAG to impact on the interaction between BIN2-YFP and Myc-BASL. Bottom, results. wt, BSL1-FLAG; mut, BSL1^D584N^-FLAG. See also Figure S4.

Our finding that BSL1 itself associates with the polarity complex (Figure 2) raises the possibility that BSL1 modulates BIN2 partitioning by actively displacing BIN2 from the polarity complex. To test this proposal, we first analyzed the ability of BSL1 to affect the stability of BIN2-BASL interactions *in vitro* and *in planta* (Figure 4B and S4E-S4H). The co-IP results show that wild-type BSL1, but not the catalytically inactive BSL1^D584N^, destabilizes BIN2-BASL interactions (Figure 4B). Next, we analyzed the ability of BSL1 to destabilize BIN2-BASL interactions *in planta* using BiFC assays (Figures S4E-S4G). Results show that BSL1, but not the catalytically inactive BSL1^D584N^, destabilizes BIN2-BASL interactions at the cell membrane in plants (Figures S4E–S4G). Furthermore, the ability of BSL1 to destabilize BIN2-BASL interactions in plants is specific, as BSL1 does not destabilize interactions between BIN2 and the scaffold protein POLAR (Houbaert et al., 2018; Pillitteri et al., 2011) (Figures S4I-S4K). The results indicate that BSL1 destabilizes BIN2-BASL interactions both *in vitro* and at the cell membrane in plants. We conclude that BSL proteins BSL1 modulate BIN2 partitioning in stomatal-lineage cells by actively displacing BIN2 from the polarity complex.

### Association of BSL1 with the Polarity Complex Activates YDA MAPK signaling

We previously showed YDA’s association with the polarity complex, like that of BSL proteins, requires interactions with BASL (Zhang et al., 2015). The association of both YDA and BSL with the polarity complex through interaction with BASL raises the possibility that YDA and BSL interact with one another. To test this proposal, we measured the ability of BSL1 to interact with YDA both *in vitro* and *in planta* (Figures 5A and S5A-S5C). Results of pull-down and co-IP assays performed *in vitro* using fusion proteins purified from *E. coli* and from leaf extracts of *N. benthamiana,* respectively, show BSL1 directly interacts with YDA (Figures 5A and S5A). Results of BiFC assays show that BSL1 interacts with YDA at the cell membrane in *N. benthamiana* leaf epidermal cells (Figure S5C). Furthermore, results of *in vivo* co-IP assays analyzing the association of fusion proteins co-expressed in *Arabidopsis* further demonstrate that BSL1 interacts with YDA (Figure S5B). Thus, we conclude that YDA directly interacts with BSL1 both *in vitro* and *in vivo*.

**Figure 5.**
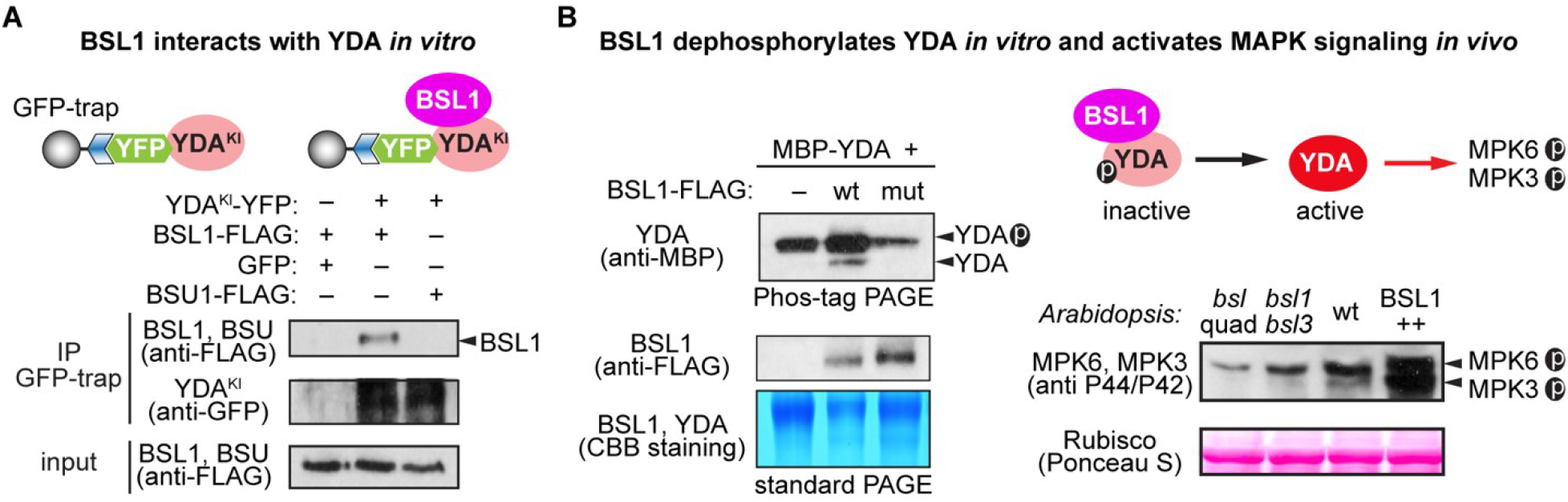
Association of BSL1 with the Polarity Complex Activates YDA and MAPK Signaling. (A) *In vitro* co-IP experiments to show BSL1 interacts with YDA. Top, schematic of co-IP assay used to monitor the interaction between BSL1-FLAG and YDA^KI^-YFP (kinase inactive to avoid overexpression-triggered cell death). Both protein fusions were overexpressed in *N. benthamiana* leaves and purified from cell extracts. Bottom, results of co-IP. (B) BSL1 dephosphorylates YDA and activates MAPK signaling. Left, *in vitro* kinase assays. Reactions were analyzed using Phos-tag PAGE (which slows the migration of phosphorylated proteins). Proteins were visualized by immunoblotting (top and middle) or CBB staining (bottom). Right, Schematic shows BSL1-mediated dephosphorylation of YDA promotes MAPK signaling *in vivo*. Blots show MPK3/6 activity levels in 3-dpg *Arabidopsis* seedlings are detected by anti-phospho-p42/p44 (top) and protein loading is shown by Ponceau S staining (bottom). See also Figure S5.

The activity of YDA is sensitive to its phosphorylation state. In particular, phosphorylation of YDA inhibits its protein kinase activity (Kim et al., 2012). Thus, the ability of BSL protein phosphatases to directly interact with YDA raises the possibility that BSL proteins directly modulate YDA activity by altering its phosphorylation state. To test this proposal, we first monitored the ability of BSL1 to modulate the phosphorylation state of YDA *in vitro* (Figure 5B, left). We incubated a recombinant MBP-YDA fusion protein with ATP (allowing auto-phosphorylation) in the presence or absence of BSL1-FLAG and visualized the phosphorylation state of MBP-YDA using Phos-tag gel electrophoresis (Figure 5B, left). In reactions containing wild-type BSL1-FLAG, but not the catalytically inactive BSL1^D584N^, we detect a faster migrating band corresponding to removal of one or more phosphate groups from MBP-YDA (Figure 5B, left). The results indicate that BSL1 dephosphorylates YDA *in vitro*. Next, we measured the ability of BSL proteins to modulate YDA-dependent MAPK signaling *in vivo* (Figure 5B, right). To do this, we monitored the phosphorylation state of two MAPKs (MPK3 and MPK6) whose activities are regulated by YDA in stomatal development (Wang et al., 2007a). Compared with levels of phosphorylated/activated MPK3 and MPK6 observed in wild-type plants, levels of phosphorylated/activated MPK3 and MPK6 are reduced in plants lacking BSL1 and BSL3 (*bsl1;bsl3*) and in plants lacking all four BSL proteins (*bsl*-quad) (Figure 5B, right). In contrast, levels of phosphorylated MPK3 and MPK6 are elevated in plants in which BSL1 is overexpressed (BSL1 ++) (Figure 5B, right). The results indicate BSL1 stimulates YDA activity *in vivo*. We conclude that the association of BSL with the polarity complex in stomatal-lineage cells activates YDA MAPK signaling.

### BSL is Required for Stomatal Development in *Arabidopsis*

The results in Figures 2 to 5 show that BSL proteins, when analyzed biochemically and at the cellular level, perform the critical functions required to alter the composition and activity of the polarity complex in the MMC *vs*. the SLGC. Accordingly, the results in Figures 2 to 5 strongly suggest BSL proteins play a key role in establishing cell-fate asymmetry during stomatal ACD in *Arabidopsis*. To directly test the role of BSL proteins in *Arabidopsis* stomatal development, we compared the generation of stomatal-lineage cells in the epidermis of wild-type plants, plants in which BSL1 was overexpressed in stomatal-lineage cells (BSL1++), and plants containing loss-of-function *bsl* alleles (Figures 6A, S6A and S6B). Results show that overexpression of BSL1 using the stomatal lineage-specific *TOO MANY MOUTHS* (*TMM*) promoter (Nadeau and Sack, 2002) suppressed formation of stomatal-lineage cells and gave rise to a leaf epidermis fully devoid of guard cells (Figures 6A and S6D). In contrast, analysis of single, double, and triple loss-of-function *bsl* mutants shows that the absence of BSL proteins leads to an increase in the percentage of stomatal-lineage cells compared with wild-type plants (Figures 6A, S6A-S6C). The results establish BSL proteins play a critical role in *Arabidopsis* stomatal development.

**Figure 6.**
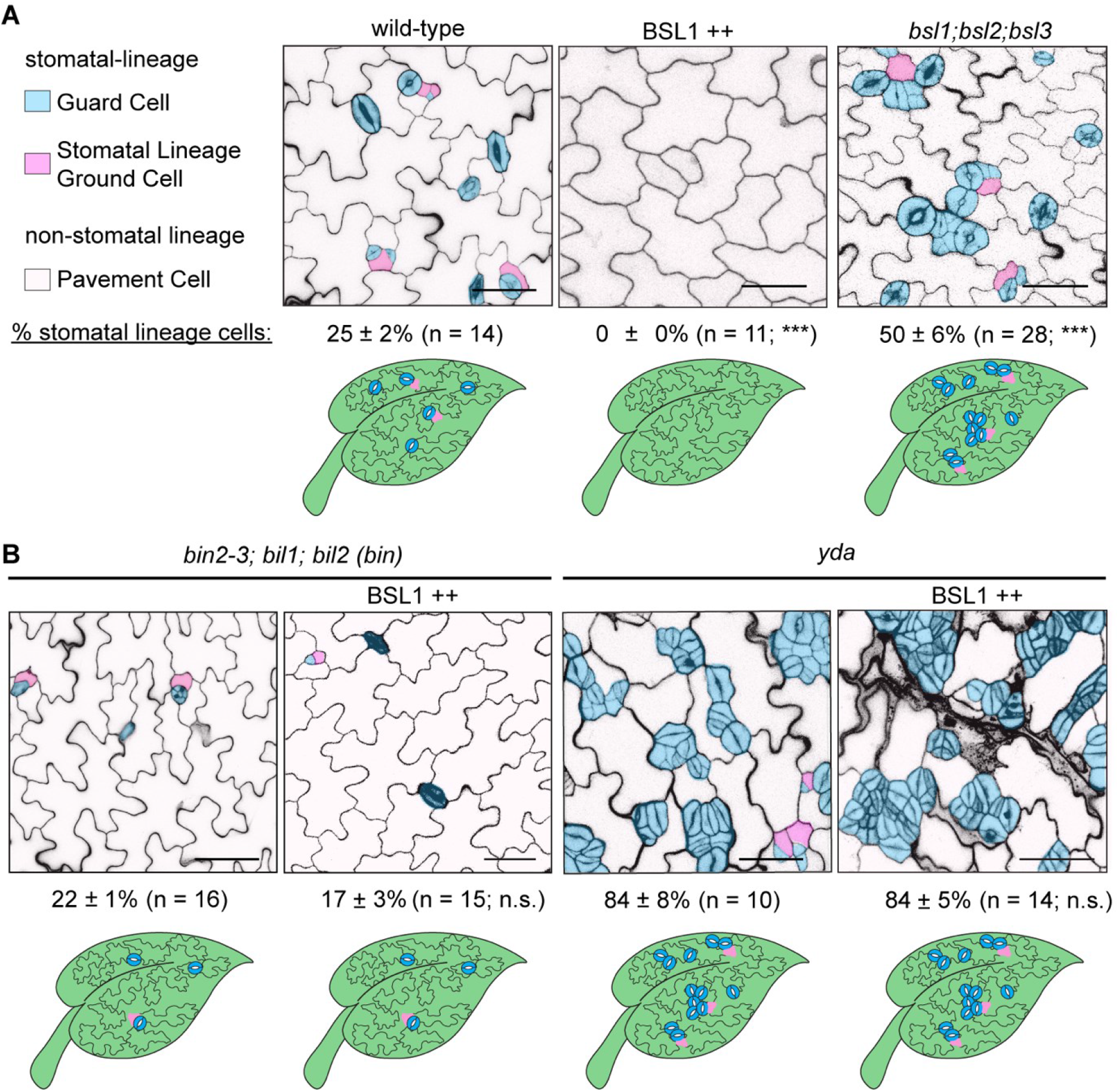
BSL Requires BIN2 and YDA to Regulate Stomatal Development in *Arabidopsis*. (A) BSL proteins function to suppress stomatal production. Five-day adaxial cotyledon epidermis of the designated genotypes were examined. BSL1 ++, overexpression of BSL1 driven by the stomatal lineage *TMM* promoter. (B) BSL requires BIN2 and YDA to regulate stomatal development. Overexpression of BSL1 (BSL1 ++) were introduced in the loss-of-function *bin* (left) or *yda* (right) mutants. (A-B) Top row: confocal images show stomatal phenotype. Cell outlines were visualized with Propidium Iodide (PI) staining. Images were captured by the confocal microscope and converted to black/white. Stomatal lineage cells were manually traced and highlighted by different shadings. Blue, stomatal guard cells; pink, SLGCs; light pink, pavement cells. Scale bars, 40 μm. Middle row: percentage of stomatal lineage cells (guard cells + SLGCs) for the corresponding genotype shown above. *n*, number of cotyledons counted. One-way ANOVA followed by Tukey’s post hoc test were used to compare with the wild type (A) or the respective mutant background (B). Data are mean ± SD (standard deviation). n.s., not significant. *** P < 0.0001. Bottom row: cartoons depict the corresponding stomatal phenotype shown above. Blue, stomatal guard cells; pink, stomatal lineage cells, puzzle shapes, pavement cells. See also Figure S6.

### BSL Function in *Arabidopsis* Stomatal Development Requires BIN2 and YDA

We next determined whether BSL proteins role in stomatal development occurs as a consequence of its ability to modulate the activities of BIN2 and YDA (Figures 6B and S6E-S6G). We first examined effects of altering BSL activity in the context of a loss-of-function *bin* mutant (carrying mutations in *BIN2* and two closely related genes *BIL1* and *BIL2*), which produces fewer stomatal-lineage cells in the leaf epidermis than wild-type plants (Figure 6B). We find that formation of stomatal-lineage cells in the loss-of-function *bin* mutant is unaffected by overexpression of BSL1 (Figures 6B and S6G). Thus, the ability of BSL proteins to modulate *Arabidopsis* stomatal development requires BIN2. Next, we examined effects of altering BSL activity in the context of a loss-of-function *yda* mutant, which overproduces stomatal guard cells, or in the context of a constitutively active *yda* mutant, which does not produce stomata (Bergmann et al., 2004)(Figures 6B, S6E and S6F). Formation of stomatal-lineage cells in plants containing either the loss-of-function *yda* mutant or plants containing a constitutively active *yda* allele is unaffected by overexpression of BSL1 or the loss-of-function *bsl-quad*, respectively (Figures 6B, S6E, and S6F). Thus, the ability of BSL proteins to modulate *Arabidopsis* stomatal development requires YDA. Taken together, the results establish that ability of BSL proteins to modulate stomatal development requires the presence of both BIN2 and YDA. We conclude that BSL’s role in stomatal development occurs as a consequence of its ability to modulate the activities of BIN2 and YDA in stomatal-lineage cells.

We also examined whether signaling components upstream of the YDA MAPK cascade are required for BSL function in stomatal development. Thus, we examined effects of BSL1 overexpression in the context of plants containing loss-of function mutations in the receptor-like protein TMM (Nadeau and Sack, 2002), in members of the ERECTA family of receptor-like kinases (Shpak et al., 2005), or in members of the SERK family of receptor-like kinases (Meng et al., 2015). We find that the effects of altering BSL activity on stomatal-lineage cell formation in these mutant backgrounds is similar to that observed in wild-type plants (Figures S6H-S6K). Thus, in contrast to YDA and BIN2, the membrane receptors TMM, ERECTA, and SERK are not required for BSL to function in *Arabidopsis* stomatal development.

## Discussion

### Polarization of BSL Proteins Provides a Spatiotemporal Molecular Switch for Stomatal ACD in *Arabidopsis*

We have identified BSL protein phosphatases as key determinants of stomatal ACD in *Arabidopsis*. Based on our findings, we propose a new mechanistic model for stomatal ACD that incorporates the critical function of BSL proteins in establishing cell-fate asymmetry (Figure 7). Prior work had shown that differences in the composition of a membrane-associated polarity complex in the stomatal ACD mother cell (MMC) *vs*. the SLGC differentiating daughter cell are required for ACD (Houbaert et al., 2018; Zhang et al., 2015). However, the mechanism by which the composition of the polarity complex changes in stomatal ACD had not been established. We have filled this knowledge gap by demonstrating that changes in the composition of the polarity occur as a consequence of polarization of BSL proteins upon the commitment of the MMC to cell division (Figure 7).

**Figure 7.**
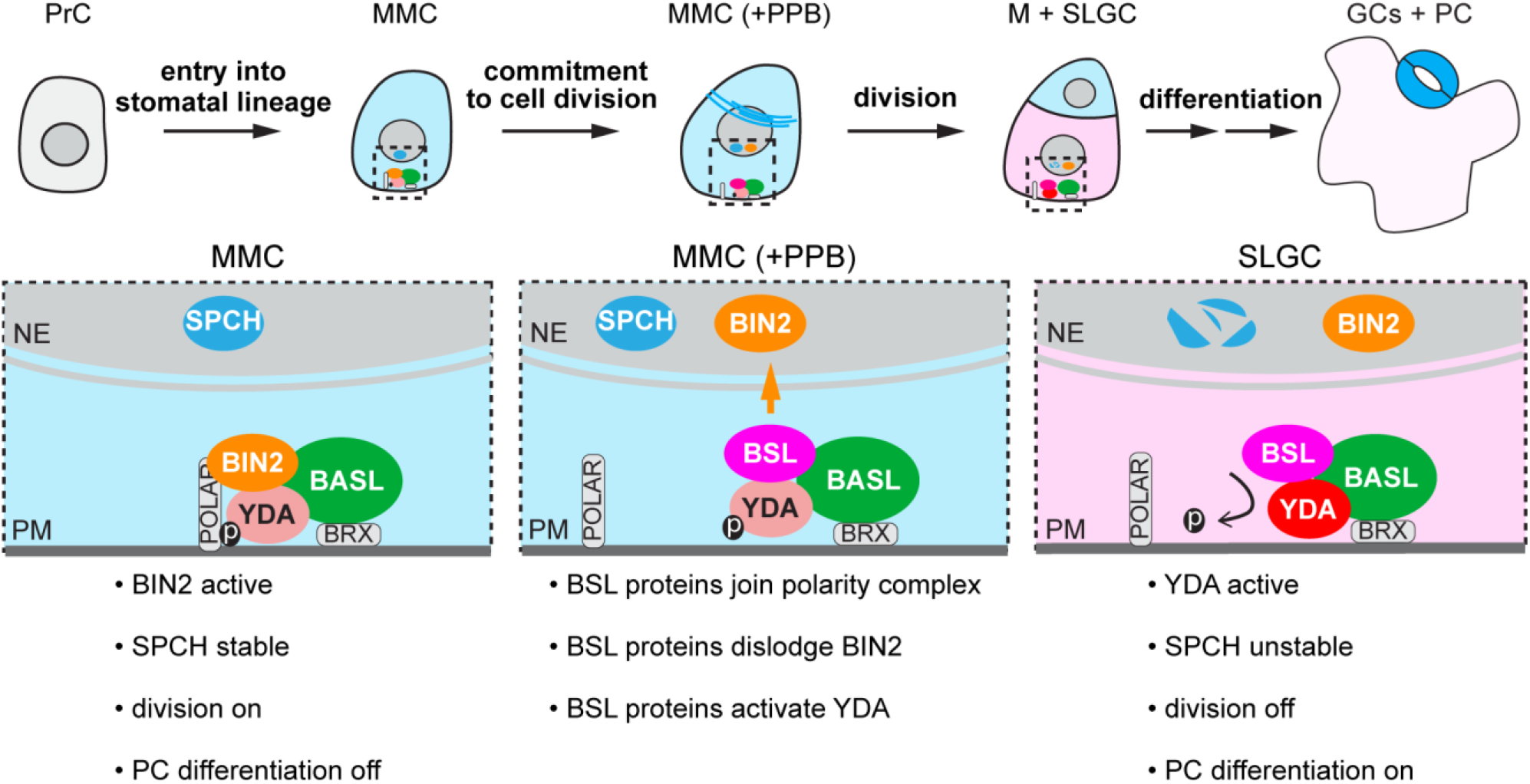
BSL Proteins Function as a Spatiotemporal Molecular Switch Enabling Stomatal ACD. Graphics on top show progressive stages of stomatal ACD in *Arabidopsis*. Dotted rectangles represent the regions enlarged at the bottom containing the polarity complex in MMC (left), PPB-containing MMC (middle) and SLGC (right). Light blue, stomatal fate; pink, non-stomatal fate. In the MMC, the BIN2 GSK3-like kinases associate with the BASL polarity complex to enrich at the cell membrane, where BIN2 suppresses the MAPKK Kinase YDA and MAPK signaling. Therefore, SPCH activity is maintained at high levels to promote cell division. The association of BSL proteins with the BASL polarity complex coincides with the formation of the PPB (blue lines in MMC), indicating the commitment of MMC to cell division. Polarized BSL is inherited by the SLGC, in which BSL displaces BIN2 from the cell membrane and activates YDA MAPK signaling, leading to strong suppression of SPCH and pavement cell differentiation. Thus, polarized BSL functions as a spatiotemporal molecular switch to establish a “kinase-based signaling asymmetry” enabling cell-fate asymmetry in the two daughter cells.

In our model, polarization of BSL proteins in the MMC establishes a “kinase-based signaling asymmetry” enabling the production of daughter cells with distinct cell-division potential and cell-fate specification. The signaling asymmetry that occurs as a consequence of BSL polarization suppresses the division of the SLGC daughter but not the meristemoid (M) daughter. Furthermore, BSL-mediated suppression of cell division allows the SLGC to differentiate into a pavement cell while the meristemoid undergoes subsequent rounds of cell division before differentiating into a guard cell. Our model predicts that plants lacking BSL activity will be unable to suppress cell division in stomatal-lineage cells, resulting in an increase in guard cell production. Consistent with this prediction, plants carrying loss-of-function *bsl* alleles exhibit stomatal over-proliferation (Figure 6). Our model further predicts that increasing BSL activity in plants will suppress cell division, resulting in a decrease in guard cell production. Consistent with this prediction, plants in which BSL1 is overexpressed do not generate stomata (Figure 6).

BSL polarization in the MMC coincides with the commitment of cells to division. Thus, BSL polarization is precisely timed to ensure the association of BSL with the polarity complex does not interfere with cell division in the MMC. Furthermore, recruitment of BSL proteins to the polarity complex at the onset of MMC cell division provides time for BSL to establish strong suppression of cell division in the SLGC. In addition, BSL polarization sequesters BSL at the cell membrane to establish the kinase-based signaling asymmetry required to suppress cell division and facilitate cell-fate specification in the SLGC but not the meristemoid. Accordingly, BSL polarization in the MMC provides a spatiotemporal molecular switch that drives cell-fate asymmetry by inhibiting cell division and activating cell-fate differentiation in the differentiating daughter cell.

### Connections Between BSL Polarization and Cell-Cycle Regulation

A key objective for future work will be to identify factors that trigger BSL polarization in the MMC. Results presented in Figure 3 show that the timing of BSL polarization in the MMC is highly correlated with the formation of the PPB, i.e., the landmark structure for cell commitment to mitosis in plants (Rasmussen et al., 2011). The demonstration that BSL polarization coincides with the initiation of mitosis in ACD progenitor MMCs suggests BSL polarization and cell-cycle regulation are connected. Studies of ACD in other eukaryotes have identified several cell-cycle regulators that impinge on the ACD machinery to promote asymmetric protein localization (e.g., the Aurora and Polo kinases, cyclins and cyclin-dependent kinases, CDKs) (Moran et al., 2019; Reich et al., 2019; Wang et al., 2007b; Witte et al., 2017). Thus, signaling components that regulate the cell cycle in *Arabidopsis* may provide temporal cues that trigger BSL polarization in the MMC. Candidate regulators that have the potential to carry out this function include the Aurora kinases, which regulate formative cell division and patterning in lateral roots (Van Damme et al., 2011), and A- and B-type CDKs, which promote the G2-M transition in plant mitosis (Vandepoele et al., 2002) and stomatal development (Boudolf et al., 2004; Yang et al., 2019). Alternatively, factors involved in the assembly of the PPB may provide the temporal cue that triggers BSL polarization. Candidate factors that have the potential to carry out this function in *Arabidopsis* include members of a “TTP” protein complex that is required for PPB assembly (Schaefer et al., 2017; Spinner et al., 2013).

### Polarization of BSL During Stomatal Development May Be Triggered by an ACD “Checkpoint”

Association of BSL proteins with the polarity complex occurs in response to a cellular mechanism that is responsive to the developmental state of the cell. We propose that a “PPB orientation checkpoint (POC),” which monitors the establishment of division-plane asymmetry, functions alongside canonical cell-cycle checkpoints to ensure the fidelity of plant ACD. According to our proposal, the POC functions, at least in part, by inhibiting BSL polarization in the MMC before the PPB has been correctly placed. Unlike animal cell division, plant cell division does not involve centrosome formation. Instead, plant cell division involves formation of spindles with axis perpendicular to the plane defined by the PPB. Thus, the POC we propose occurs in plant ACD would be functionally analogous to the “spindle position checkpoint” (SPOC) described in budding yeast (Lew and Burke, 2003) and the “centrosome orientation checkpoint” (COC) described in *Drosophila* germ lines (Venkei and Yamashita, 2015).

In symmetric cell divisions, cell-cycle checkpoints monitor the of the major cellular events (cell size, DNA integrity, chromosome replication and segregation) to ensure the fidelity of the cell cycle (Hartwell and Weinert, 1989). The absence of these checkpoints has deleterious consequences for the development of cells and organs. ACD requires additional checkpoints to ensure the position and orientation of the spindle will result in the production of asymmetric daughter cells (Lew and Burke, 2003; Venkei and Yamashita, 2018). Accordingly, the absence of ACD checkpoints can result in stem cell over-proliferation due to symmetric self-renewal or stem-cell depletion due to symmetric differentiation of daughter cells. In this regard, stem cell over-proliferation and stem cell depletion is precisely what we observed in plants in which BSL activity has been perturbed.

### Signaling Complexity Underlies Developmental Plasticity in Plants

The sessile lifestyle of plants requires plant growth and developmental processes to adapt to changing environments. Thus, signaling pathways controlling growth and development in plants exhibit greater flexibility, i.e., “developmental plasticity,” compared with pathways controlling growth and development in mammalian systems. In metazoan stem cell ACD, polarization of the conserved PAR proteins in the mother cell sequesters cell-fate determinants, thus ensuring they are inherited by only one of the two daughter cells (Goldstein and Macara, 2007; Neumuller and Knoblich, 2009). In *Arabidopsis* stomatal stem cell ACD, polarization of BASL in the mother cell provides a platform that enables “kinase-based signaling asymmetry” in the daughter cells. Both the “protein sequestration” mechanism used in metazoans and the “signaling asymmetry” mechanism used in plants enable the establishment of cell-fate asymmetry with high fidelity. In stomatal ACD, the major cell-fate determinant (SPCH) and its regulators (BIN2, YDA, BSL) are each responsive to environmental changes (Dóczi and Bögre, 2018; Farkas et al., 2007). Thus, stomatal ACD, unlike metazoan stem cell ACD, exhibits developmental plasticity that allows plants to readily modulate stomatal production in response to environment cues such as light and temperature (Gudesblat et al., 2012; Lau et al., 2018). Additionally, the functional connections between BIN2, YDA, BSL and SPCH may integrate and coordinate basic core cellular events, e.g. cell size and cell shape (Kim and Wang, 2010; Lukowitz et al., 2004), to maintain the fine balance between cell division and cell differentiation.

## SUPPLEMENTAL INFORMATION

- Materials and Methods
- Figures S1-S6
- Tables S1

## ACKNOWLEDGEMENTS

We thank the ABRC stock center for providing T-DNA insertional seeds, the Biological Mass Spectrometry Facility of Robert Wood Johnson Medical School for performing mass spectrometry analysis, and Jared Winkelman for assistance with data analysis. This research was supported by NIH grants GM109080 and GM131827 to J.D., GM066258 to Z-Y.W., and GM118059 to B.E.N.

## AUTHOR CONTRIBUTIONS

X.G. and J.D designed the research. X.G. performed most of the experiments. C.H.P. assisted with mutant analysis. X.G., C.H.P, Z-Y.W., B.E.N. and J.D. wrote the manuscript.

## DECLARATION OF INTERESTES

The authors declare no competing interests.

