## Supplemental Info for "A Spatiotemporal Molecular Switch Governs Plant Asymmetric Cell Division"

**This file includes:**

Materials and Methods
Figures S1 to S6
Table S1

### MATERIALS AND METHODS

#### Plant materials

The *Arabidopsis thaliana* ecotype Columbia (Col-0) was used as the wild-type. All mutants are in the Col-0 background except for *bin2-3;bil1;bil2* in the Wassilewskija (Ws-0) background. The *BSLf* T-DNA insertional mutants were obtained from the *Arabidopsis* Biological Resource Center (ABRC), including *bsl1-1* (SALK_051383), *bsl1-2* (SALK_147279), *bsl2-1* (SALK_055335), *bsl2-2* (WiscDsLox245G08), *bsl3-1* (SALK_071689), *bsl3-2* (SALK_072437), *bsu1-1* (SALK_030721), and *bsu1-2* (SAIL_101_H03). The null alleles, *bsl1-1* (SALK_051383), *bsl2-1* (SALK_055335), *bsl3-2* (SALK_072437) and *bsu1-1* (SALK_030721), were used for generating high order mutants and phenotypic analysis. The bsl family mutants were confirmed by PCR-based genotyping. Mutants and transgenic *Arabidopsis* lines reported previously were: *basl-2*, *pBASL*::GFP-BASL*;basl-2, and p35S*::GFP-BASL*;basl-2* (Dong et al., 2009), *pSPCH*::SPCH-CFP*;spch-3* (Lau et al., 2014), *yda-3* (Salk_105078) (Zhang et al., 2015), *bsl*-quad (null alleles of *bsu1*, *bsl1* combined with RNAi-mediated silencing of *BSL2* and *BSL3*) and *bin2-3;bil1;bil2* (Kim et al., 2012), *tmm* (Nadeau and Sack, 2002), *er;erl1;erl2* (Shpak et al., 2005), *serk-q* (Meng et al., 2015) and *p35S*::mCherry-TUA5 (Gutierrez et al., 2009). Primers for genotyping, reverse transcription (RT)-PCR and quantitative real-time PCR are listed in Supplemental Table 1.

#### Growth conditions

In general, *Arabidopsis* seeds were surface sterilized with 10% bleach and grown on half-strength Murashige and Skoog (MS) basal medium plates or in soil with 16-hr light/8-hr dark cycles at 22 °C. Wild-type *N. benthamiana* plants were grown under 14-hr light and 10-hr darkness at 25 °C.

#### Plasmids construction

The Gateway cloning technology (Invitrogen) was used for most DNA manipulations unless otherwise specified. For molecular cloning, the coding DNA sequences (CDS) and genomic DNA fragments (gDNA) of *BSL1*, *BSL2*, *BSU1* and *BIN2* were cloned into pENTR/D/TOPO vectors (Invitrogen) and for *BSL3*, the CDS was cloned into pENTR/D/TOPO. The phosphatase-dead BSL1$^{D584N}$ was obtained by site-directed mutagenesis of *BSL1* CDS in pENTR/D/TOPO by QuikChange II Site-Directed Mutagenesis Kit (Agilent). The kinase inactive version of YDA (YDA$^{KI}$ containing the site mutation K429R) (Lampard et al., 2009) was cloned into pENTR/D/TOPO vectors. The *BASL* and *TMM* promoter sequences can be found in (Dong et al., 2009) and (Nadeau and Sack, 2002) and were subcloned into pDONR-P4-P1R, respectively (courtesy of Dr. Diego Wengier). Double LR recombination reactions using LR Clonase II (Invitrogen) were performed to integrate pENTR/D containing CDS or gDNA of the gene-of-interest and pDONR-promoter into the R4pGWB vectors (Nakagawa et al., 2008). To generate constructs for protein localization, the promoter regions of *BSL1*, *BSL2*, *BSL3* and *BSU1* were isolated by PCR using the primers specified in Supplemental Table 1 and inserted into the corresponding pENTR/D-CDS plasmids. For BIN2 protein fusion, the genomic fragment containing a 2.2 kb promoter and the genomic coding region of *BIN2* was amplified and cloned into pENTR/D/TOPO, and then recombined into pHGY (Kubo et al., 2005). For transient protein expression in *N. benthamiana*, the pENTR/D vectors containing the coding sequences of *BSL1*, *BSL2*, *BSL3, BSU1, BIN2, BASL, or POLAR* were recombined into the pH35GC/Y (Kubo et al., 2005) or pGWB to generate *p35S*::BSLf-CFP, *p35S*::BSLf-FLAG, *p35S*::BIN2-YFP, *p35S*::4xMyc-BASL, or *p35S*::4xMyc-POLAR. The pXNGW and/or pXCGW vectors were used for recombination reactions to generate the BiFC constructs. The resulted binary vectors mentioned above were confirmed by restriction enzyme digestion and DNA sequencing, then transferred into *Agrobacterium tumefaciens* strains GV3101 for *Arabidopsis* transformation and/or *N. benthamiana* leaf infiltration.

**Confocal imaging and image processing**

Confocal images were acquired by a Leica TCS SP5 II microscope. The excitation/emission spectra for various fluorescent proteins are: CFP, 458 nm/480–500 nm; GFP, 488 nm/501–528 nm; YFP, 514 nm/520–540 nm; mCherry, 543 nm/600–620 nm, RFP, 594 nm/600–620 nm, and propidium iodide (PI), 594 nm/591–636 nm. Images were taken from similar central areas in the adaxial side of developing cotyledons of *Arabidopsis* seedlings or *N. benthamiana* leaves. Cells outlines in *Arabidopsis* were visualized by propidium iodide (PI, Invitrogen) staining. All imaging processing was performed with Fiji (Image J) software (http://fiji.sc/Fiji). Whenever possible, z-stacked images were obtained. Quantifications and statistical analyses were performed using Fiji and GraphPad Prism 5.1, respectively.

**Recombinant protein expression, purification and in vitro pull-down assay**

To express recombinant proteins in *E. coli*, the coding region of *BASL* was cloned into pET28a vector to generate His–tagged BASL. The coding region of *BSL1* was cloned into pMAL-c2x and pGEX-4T-1 vector to generate MBP–tagged BSL1 and GST-tagged BSL1, respectively. The coding region of *BIN2* was cloned into pGEX-4T-1 vector to generate GST-tagged BIN2. The coding regions of YDA was cloned into pMAL-c2x vector to generate MBP–tagged YDA. The primer sequences are listed in Supplemental Table 1.

All constructs were transformed into *E. coli* (BL21 strain) and recombinant protein expression was induced by IPTG (0.5 mM) at 16 °C for 16 hrs. Bacterial cells were harvested and lysed by sonication in lysis buffer (50 mM Tris-HCl pH7.4, 150 mM NaCl, 0.5 mM EDTA and 1 mM PMSF). GST-tagged proteins were purified with Pierce™ Glutathione Superflow Agarose (Thermo Scientific™), MBP-tagged proteins were purified with Amylose Resin (NEB), and His-tagged proteins were purified with Ni-NTA Agarose (Qiagen) according to the manufacturer's instructions.

Purified proteins were used for *in vitro* pull-down assays. GST or GST-BSL1 proteins were immobilized on Pierce™ Glutathione Superflow Agarose, which were then incubated with equal amount of purified MBP-tagged YDA at 4 °C for 3 hrs. The beads were washed three times with the washing buffer (50 mM Tris-HCl pH 7.4, 150 mM NaCl, 0.5 mM EDTA, 0.5% TritonX-100). Proteins bound to the beads were mixed with 5 × SDS sample buffer and boiled for 5 mins. Samples were separated by 10% sodium dodecyl sulphate-polyacrylamide gel electrophoresis (SDS–PAGE) and transferred onto a polyvinylidene fluoride membrane (Bio-Rad). Protein were analyzed by Immunoblot with anti-MBP (Anti-MBP Monoclonal Antibody, NEB, E8032S,1:5000) or anti-GST (GST (91G1) Rabbit mAb #2625, Cell Signaling Technology, 1:1000) antibodies, respectively.

*In vitro* quantitative pull-down assays (Figure S6D) were performed as previously described (Houbaert et al., 2018) to examine different binding affinities of BIN2-BASL in the presence of BSL1. All testing proteins (MBP/MBP-BSL1, GST-BIN2, and His-BASL) were purified beforehand. To assay the interaction levels of BIN2-BASL, Ni-NTA Agarose was first used to immobilize His-BASL, the mixture was then split into 4 tubes with equal amount. For each reaction, same amount of GST-BIN2 with increasing amount of MBP-BSL1 or MBP (negative control) (1x, 5x, and 10x concentrated) were added into same amount of His-BASL and incubated at 4 °C for 3 hrs followed by washing buffer, SDS-PAGE and Immunoblotting. The BIN2-BASL interaction strength were evaluated by the amount of GST-BIN2 being pulled down with His-BASL. His-BASL was detected by Anti-His (Antibody, #2365, Cell Signaling Technology, 1:1000). To quantify protein amount on Immunoblots, the band intensities (3 times of repeats) were measured by Fiji. To generate the histograms, relative amount of target proteins (GST-BIN2) were first normalized to the BASL band intensity and the amount of GST-BIN2 in the control reaction (His-BASL + GST-BIN2 without BSL1) was defined as 1.

**In vitro kinase assay**

To evaluate *in vitro* phosphorylation levels of YDA, 0.5 μg of purified MBP-YDA (EDTA free) with or without BSL1-FLAG/BSL1$^{D584N}$-FLAG fusion proteins purified from *N. benthamiana* leaf were incubated in 30 μl reaction buffer (5 mM HEPES, 10 mM $MgCl_2$, 10 mM $MnCl_2$, 1 mM DTT and 10 μM cold ATP) at 30 °C for 30 mins. Reactions were stopped by adding 6 μl of 5 × SDS sample buffer. Protein samples were analyzed on a Phos-tag acrylamide gel (50 μM, Wako Chemicals) that slows down the migration of phosphorylated proteins and separates YDA with different phosphorylation levels, Protein loading were performed by Immunoblot analysis with anti-MBP antibody (Anti-MBP Monoclonal Antibody, NEB, E8032S,1:5000).

**Transient expression in *Nicotiana benthamiana* and imaging analysis**

*Agrobacterium* strains GV3101 harboring the constructs-of-interest in 10 ml of LB medium with appropriate antibiotics were cultured overnight. Bacterial cells were collected at 4500 rpm for 10 mins and resuspended in 10 mM $MgCl_2$. Cell culture and p19 were mixed to reach an $OD_{600}$ = 0.5 for each line and co-infiltrated into the 4-week-old *N. benthamiana* abaxial leaves as described previously (Zhang et al., 2015). Co-infiltrated leaves were checked by confocal microscopy 3-4 days post infiltration. To quantify the polarity degree of YFP in *N. benthamiana* epidermal cells, three independent replicate experiments were performed and 26–28 representative cells for each combination were scored. Absolute fluorescence intensity values were measured by Fiji and protein polarization were calculated as described in (Zhang et al., 2015).

**Co-immunoprecipitation (co-IP) assay in *Arabidopsis* and *N. benthamiana***

To test *in vivo* physical interaction between BSL1 and YDA, total cell proteins were extracted from *Arabidopsis* plants (3-dpg seedlings) co-expressing *pTMM*::*BSL1-FLAG and pBASL*::*10xmyc-YDA$^{KI}$*. To perform the experiments, plant tissues were ground up in liquid nitrogen, and then mixed with the protein extraction buffer (100 mM Tris-HCl pH 7.5, 5 mM EDTA, 5 mM EGTA, 1 mM $Na_3VO_4$, 10 mM NaF, 50 mM b-glycerophosphate, 10 mM DTT, 1 mM phenylmethylsulfonyl fluoride, 5% (v/v) glycerol, 0.5% (v/v) Triton X-100, and 1% (v/v) protease inhibitor cocktail (Sigma-Aldrich, P 9599)). Protein extracts were centrifuged at 12,000g at 4°C for 20 mins and the supernatants were incubated with Anti-FLAG M2 Affinity Gel (Sigma-Aldrich) at 4°C for 3 hrs. After incubation, the immunoprecipitated proteins with beads were washed three times with the extraction buffer, then mixed with 2 × SDS sample buffer and boiled for 5 mins. Samples were separated by 10% SDS–PAGE followed Immunoblot by corresponding primary antibodies (Monoclonal Anti-FLAG antibody produced in rabbit, F2555, Sigma-Aldrich; Myc-Tag (71D10) Rabbit mAb #2278, Cell Signaling Technology).
To test protein-protein interactions of BASL-BSL and BSL1-YDA in plant cells, total cell proteins were extracted from *N. benthamiana* leaves that transiently expressed the combinations of *p35S*::*GFP* or *p35S*::*GFP-BASL* with *p35S*::*BSL1-FLAG*, *p35S*::*BSL2-FLAG*, *p35S*::*BSL3-FLAG,* or *p35S*::*BSU1-FLAG* and the combinations of *p35S*::*YDA$^{KI}$-YFP* with *p35S*::*BSL1-FLAG* or *p35S*::*BSU1-FLAG*. To test the influence of BSL1/BSL1$^{D584N}$ (phosphatase-dead) on the interactions of BASL-BIN2 or POLAR-BIN2, total cell proteins were extracted from *N. benthamiana* leaves transiently co-expressing *p35S*::4xmyc-BASL/POLAR and *p35S*::BIN2-YFP in the presence of *p35S*::BSL1/BSL1$^{D584N}$-FLAG. *N. benthamiana* leaves were collected and ground in liquid nitrogen with the same extraction buffer described above. Protein extracts were centrifuged at 18,000g at 4 °C for 30 mins and the supernatants were incubated with GFP-Trap Agarose beads (Chromotek) at 4 °C for 3 hrs. Then, the beads were washed three times with

extraction buffer, followed by mixing with 2 × SDS sample buffer and boiling for 5 mins. Samples were separated by 10% SDS–PAGE and analyzed by corresponding primary antibodies (Monoclonal ANTI-FLAG M2 antibody produced in mouse, F3165, Sigma-Aldrich; Anti-GFP Antibody, Roche (Cat#11814460001), Myc-Tag (9B11) Mouse mAb #2276, Cell Signaling Technology).

**Immunoprecipitation–mass spectrometry (IP–MS) analysis**

Five grams of seedlings (expressing *pBASL*::GFP-BASL in *basl-2*, or *p35S*::GFP-BASL in *basl-2*, or *pBASL*::GFP in Col-0 at 3-dpg) were ground to a fine powder in liquid nitrogen. Total cell proteins were extracted with extraction buffer (100 mM Tris-HCl at pH7.5, 150 mM NaCl, 5 mM EDTA, 5 mM EGTA, 10 mM DTT, 10 mM $Na_3VO_4$, 20 mM NaF, 50 mM β-glycerophosphate, 10% glycerol, 1 mM PMSF, protease inhibitor cocktail for plant cell extracts (Sigma-Aldrich, P 9599), 1% (v/v) NP-40). The homogenates were sonicated for 10 seconds in ice bucket, then dilute with extraction buffer without NP-40 to lower NP-40 to 0.2% (v/v). Centrifuge the protein mix at 10,000g for 30 mins at 4 °C to collect supernatant. Protein concentration of the supernatant was measured by aliquoting 50 μl for the Protein Assay Reagent (BioRad). The remaining supernatant was mixed with 100 μl GFP-Trap Agarose (Chromotek) and incubated for 3 hrs on a rotating wheel at 4 °C. The beads were then collected by low speed centrifuge and washed four times by resuspending in protein extraction buffer with 0.2% (v/v) NP-40. Add 5 × SDS sample buffer into the beads and boil at 95°C for 5 mins. Protein samples were loaded on 10% SDS-PAGE but run for a short distance, followed by gel reduction, alkylation and digestion with trypsin (sequencing grade, Thermo Scientific, Cat# 90058). Digested peptides in the gel were extracted twice with 5% formic acid, 60% acetonitrile and dried under vacuum. Peptide samples were analyzed by LC-MS using Nano LC-MS/MS (Dionex Ultimate 3000 RLSC nano System) interfaced with QExactive HF (Thermo Fisher). Peptides were loaded on to a fused silica trap column Acclaim PepMap 100, 75 um x 2 cm (Thermo Fisher). After washing for 5 mins at 5 μl/min with 0.1% TFA, the trap column was brought in-line with an analytical column (Nanoease MZ peptide BEH C18, 130A, 1.7 um, 75 um x 250 mm, Waters) for LC-MS/MS. Peptides were fractionated at 300 nL/min using a segmented linear gradient 4-15% B in 30 mins (where A: 0.2% formic acid, and B: 0.16% formic acid, 80% acetonitrile), 15-25% B in 40 mins, 25-50% B in 44 mins, and 50-90% B in 11 mins.
Mass spectrometry data were obtained using a data-dependent acquisition procedure with a cyclic series of a full scan acquired in Orbitrap with resolution of 120,000 followed by MS/MS (HCD relative collision energy 27%) of the 20 most intense ions and a dynamic exclusion duration of 20 sec. The peak list of the LC-MS/MS were generated by Thermo Proteome Discoverer (v. 2.1) into MASCOT Generic Format (MGF) and searched against *Arabidopsis* (TAIR v. 10), plus a database composed of common lab contaminants using in house version of X!Tandem (GPM Furry, Craig and Beavis, 2004). Search parameters are as follows: fragment mass error: 20 ppm, parent mass error: +/- 7 ppm; fixed modification: carbamidomethylation on cysteine; flexible modifications: Oxidation on Methionine; protease specificity: trypsin (C-terminal of R/K unless followed by P), with 1 miss-cut at preliminary search and 5 miss-cut during refinement. Only spectra with loge <-2 were included in the final report. LC-MS/MS analysis was performed at the Biological Mass Spectrometry facility of Rutgers University.

**MAPK activation assay**

Total cell proteins were extracted with extraction buffer (50 mM HEPES pH 7.5, 150 mM NaCl, 5 mM EDTA, 5 mM EGTA, 10 mM DTT, 10 mM $Na_3VO_4$, 20 mM NaF, 50 mM β-glycerophosphate, 10% glycerol, 1 mM PMSF, protease inhibitor cocktail for plant cell extracts (Sigma-Aldrich, P 9599), 1% (v/v) NP-40) from 3-dpg *Arabidopsis* seedlings of Col-0, *bsl1;bsl3*, *bsl-quad*, and *pTMM*::BSL1-YFP. The extracted total cell proteins were resolved on 10% SDS–PAGE, followed by Immunoblot with the primary antibody against phosphor-p42/44 MAP kinase

(1:1000, Cell Signaling Technology). Total protein staining (Ponceau S) was used to confirm equal loading for Immunoblots.

**Yeast two-hybrid assay**

Full-length coding sequence of *BSL1*, *BSL2*, *BSL3*, or *BSU1* was cloned and inserted into the pGBKT7 vector as bait. The full-length coding sequence of *BASL* was cloned into the pGADT7 vector. Constructs used for testing the interactions were co-transformed into *Saccharomyces cerevisiae* strain AH109 using EZ-YEAST™ transformation kit (MP Bio-medicals) following the manufacturer's instructions. The positive transformants were selected on the SD/-Leu/-Trp medium. The interactions were tested on the SD/-Leu/-Trp/-His medium with appropriate concentration of 3-amino-1,2,4-triazole (3-AT). Interactions were observed after 3 days of yeast growth at 30 °C.

**Quantitative real-time PCR**

Total RNAs were extracted from 50 mg of seedlings at 3 to 5-dpg with RNeasy Plant Mini Kit (Qiagen). cDNAs were generated by the SuperScript™ III First-Strand Synthesis System (Invitrogen). Transcript levels of *BSL1*, *BSL2*, *BSL3*, *BSU1* were amplified with the primers listed in Supplemental Table 1 and the reactions were set up by SYBR™ Select Master Mix (Thermo Fisher Scientific). *Actin 2* was used as an internal control to normalize expression levels. Data are presented as mean ± SD. Quantitative real-time PCRs were performed by the Stepone real-time PCR system (Applied Biosystems).
For reverse transcription (RT)-PCR, two independent lines from each mutant were tested. The ribosomal *S18* (*RPS18*) gene was used as an internal standard for normalization of gene expression levels. The sequences of all primers used are listed in Supplemental Table 1.

**QUANTIFICATION AND STATISTICAL ANALYSIS**

**Stomatal quantification and protein polarity quantification**

In general, to examine stomatal phenotypes in development, 5-dpg (days post germination) cotyledons were stained with PI (Invitrogen) to capture images from similar central regions of adaxial cotyledons. Images were captured by the EC Plan-Neofluar (20×/0.5) lenses on a Carl Zeiss Axio Scope A1 fluorescence microscope equipped with a Progress MF CCD camera (Jenoptik). Typically, 12–20 individual seedlings were picked from each mutant or two representative T2 transgenic lines out of >12 independent transgenic events. Confocal images shown in the figures were false colored with brightness/contrast adjusted by Fiji.

To quantify stomatal phenotypes, the epidermal cells were categorized by size and shape into three groups: guard cells (pairs of kidney-shaped), stomatal lineage cells (small dividing cells, including MMCs, Ms and SLGCs), and pavement cells (puzzle-shaped epidermal cell and enlarged SLGCs with at least one obvious lobe). Quantification for stomatal index (SI: number of stomata/total number of epidermal cells) and stomatal lineage index (SLI: number of guard cell pairs + stomatal lineage cells over total number of epidermal cells) were calculated by counting cells in an area of 0.385 $mm^2$ with the cell-counter plug-in in Fiji.

To quantify protein polarization (BSL1 and BASL) in different cell types in *Arabidopsis*, confocal images were taken from adaxial cotyledons of 10-15 representative seedlings at 36-hpg, 48-hpg, and 72-hpg. Cells were subdivided into four categories based on cell morphology combined with BASL expression patterns: "protodermal cells" (PrC), small rectangular cells with nuclear-only BASL; "early MMCs", small rectangular MMCs expressing both nuclear and polarized BASL; "late MMCs", elongated/asymmetric MMCs expressing nuclear and polarized

BASL; and "SLGCs", large daughter cells after cell division completed with polarized BASL. For each category, 30-50 individual cells were selected and scored for protein polar distribution. To measure protein polarization values, ratios of high fluorescence intensity values (a) over low fluorescence intensity values (b) along equal cell periphery lengths from the same cell were collected and calculated. For the cells with relatively uniform expression of fluorescent proteins, the a and b values were collected by measuring two randomly selected cell periphery with equal length.

To quantify SPCH expression pattern, seedlings expressing SPCH-CFP in the wild-type and *bsl-quad* mutants were stained with PI and images of the adaxial side cotyledon epidermis were captured by Confocal. Numbers of CFP-positive stomatal lineage cells and total epidermal cells were counted with Fiji to obtain the CFP-positive ratios.

**Statistics and reproducibility**

All statistical analyses were conducted with GraphPad Prism 5.1 Software. To compare two normally distributed groups, unpaired two-tailed t-tests were used. For multiple comparisons between normally distributed groups, one-way ANOVA followed by Tukey's post hoc test were used. For all figures, *P < 0.05, **P < 0.005, ***P < 0.0001. To count cell types and to quantify immunoblots, the grid and image counter plug-ins in Image J were used. Numbers of repetitions and replicates for each experiment were mentioned in the legends.

### Supplemental Figures and Figure Legends

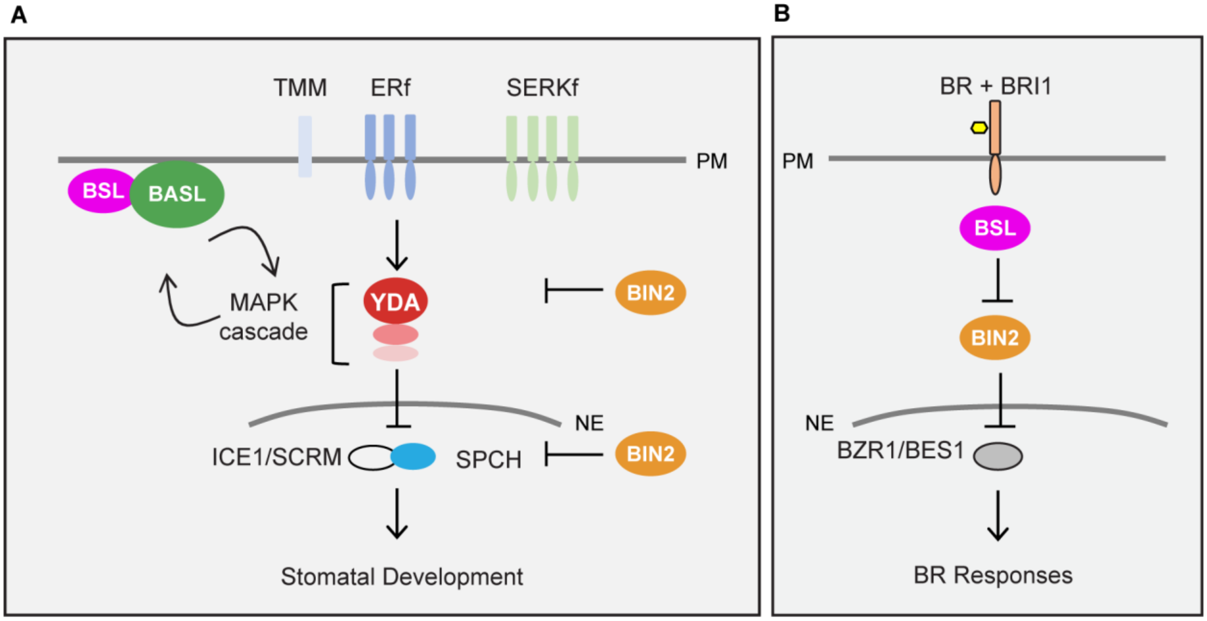


**Figure S1. Genetic Pathways Involving BSL Proteins in *Arabidopsis*, Related to Figure 1**
Schematics depicting signal transduction pathways in stomatal development (A) and Brassinosteroid (BR) signaling (B). The models are mainly based on *Arabidopsis* research. Both pathways involve BSL Ser/Thr protein phosphatases and the GSK3-like BIN2 kinase. Block lines indicate negative regulation and arrows indicate positive regulation. PM, plasma membrane; NE, nuclear envelope.

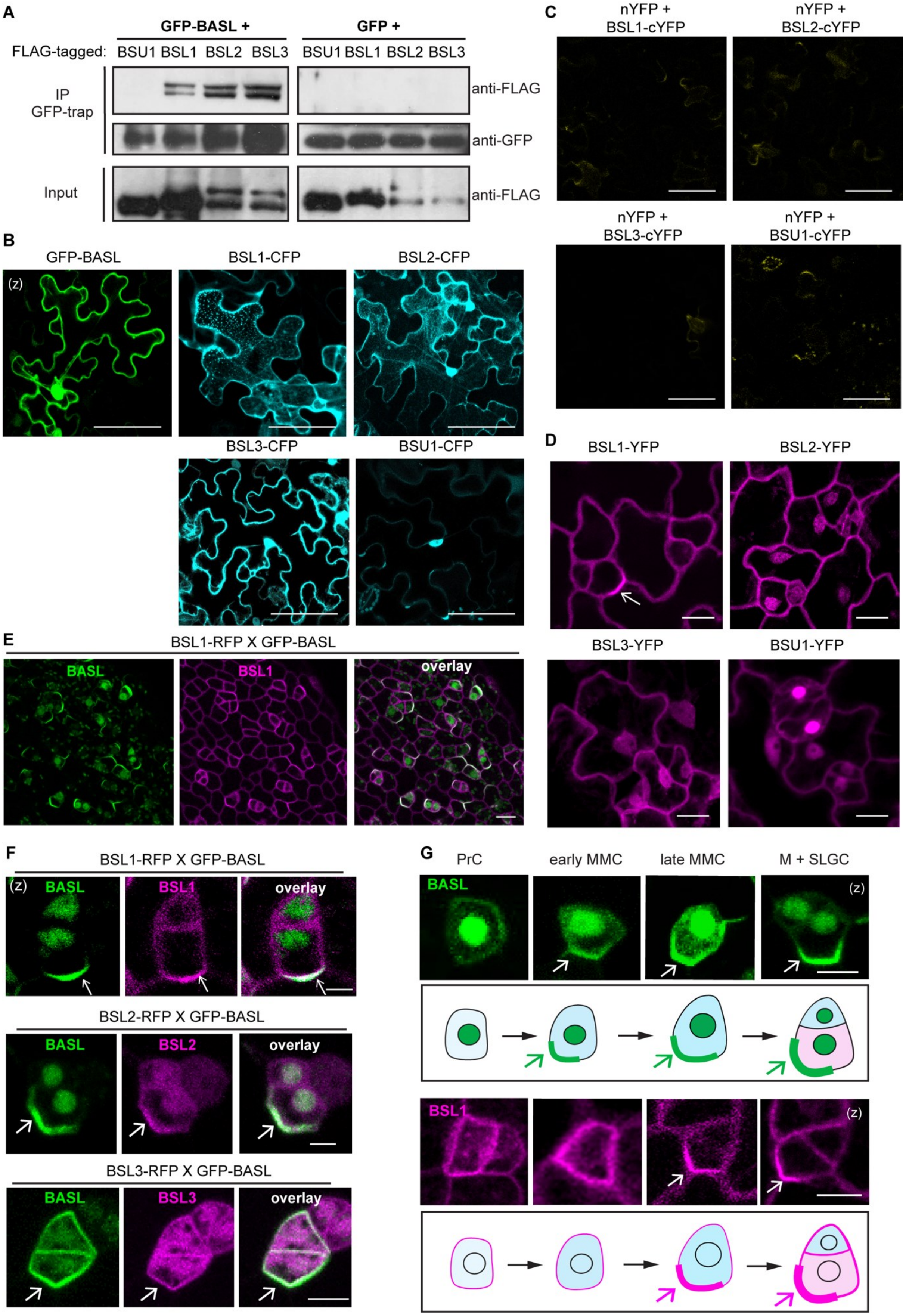
A
GFP-BASL +
GFP +
FLAG-tagged: BSU1 BSL1 BSL2 BSL3 BSU1 BSL1 BSL2 BSL3
IP
GFP-trap
Input
anti-FLAG
anti-GFP
anti-FLAG
B
GFP-BASL
BSL1-CFP
BSL2-CFP
BSL3-CFP
BSU1-CFP
(z)
C
nYFP +
BSL1-cYFP
nYFP +
BSL2-cYFP
nYFP +
BSL3-cYFP
nYFP +
BSU1-cYFP
D
BSL1-YFP
BSL2-YFP
BSL3-YFP
BSU1-YFP
E
BSL1-RFP X GFP-BASL
BASL
BSL1
overlay
F
BSL1-RFP X GFP-BASL
(z)
BASL
BSL1
overlay
BSL2-RFP X GFP-BASL
BASL
BSL2
overlay
BSL3-RFP X GFP-BASL
BASL
BSL3
overlay
G
PrC
early MMC
late MMC
M + SLGC
BASL
(z)
BSL1
(z)

**Figure S2. BSL1, BSL2 and BSL3 Physically Interact and Polarize with BASL *in vivo*, Related to Figure 2**

(A) Co-IP assays using purified fusion proteins transiently expressed in *N. benthamiana* leaf cells. Results show physical association of FLAG-tagged BSL1/2/3 proteins with GFP-BASL. GFP-BASL or GFP (control) was used as bait to bind to the GFP-Trap Agarose. Immunoprecipitated proteins were detected by anti-FLAG. Data represent results of three biological repeats.

(B) Representative confocal images show subcellular localization of indicated proteins in *N. benthamiana* leaf epidermal cells. BASL (green), nuclear and cytoplasmic; BSL1 (cyan), cell membrane and cytoplasmic puncta; BSL2 (cyan), nuclear and cytoplasmic; BSL3 (cyan), nuclear and cytoplasmic; BSU1, mainly nuclear and weakly cytoplasmic. (z), z-staked confocal images.

(C) Negative controls for the BiFC assays to test BSL-BASL interaction in *N. benthamiana* leaf epidermal cells. Recovered YFP signals suggest positive protein-protein interaction and no significant YFP signals were recovered when nYFP was coupled with BSL-cYFP. nYFP and cYFP, N- and C-terminal domain of the split YFP, respectively. Scale bars in (B-C), 50 μm.

(D) Subcellular localization of the BSL proteins (magenta) in stomatal lineage cells in 3-dpg *Arabidopsis* adaxial cotyledon epidermis. The expression of BSL1 and BSL2 were driven by their native promoters and the expression of BSL3 and BSU1 were driven by the ubiquitous *35S* promoter. Consistent with data shown in (B), BSL1 was found in the cytoplasm and close to the cell membrane. BSL2, BSL3 and BSU1 were found in the cytoplasm and nucleus. Arrow indicates polarized localization of BSL1 in stomatal lineage cells. Scale bars, 10 μm.

(E) Co-expression of GFP-BASL (green) with BSL1-RFP (magenta), both driven by the *BASL* promoter, in a true leaf of 5-dpg seedling.

(F) Representative images of co-localization of BSL1-RFP, BSL2-RFP and BSL3-RFP (magenta) with GFP-BASL (green), all driven by the *BASL* promoter. Arrows mark protein polarization at the cell membrane.

(G) Expression patterns of endogenous promoter driven GFP-BASL (green) and BSL1-YFP (magenta) in progressive cell types during stomatal development. PrC (protodermal cells), early MMC (Meristemoid Mother Cell, small and rectangular), late MMC (asymmetrically expanded and triangle), M (Meristemoid), and SLGC (Stomatal Lineage Ground Cell). Protein polarizations were indicated by arrows. Scale bars in (E-G), 5 μm.

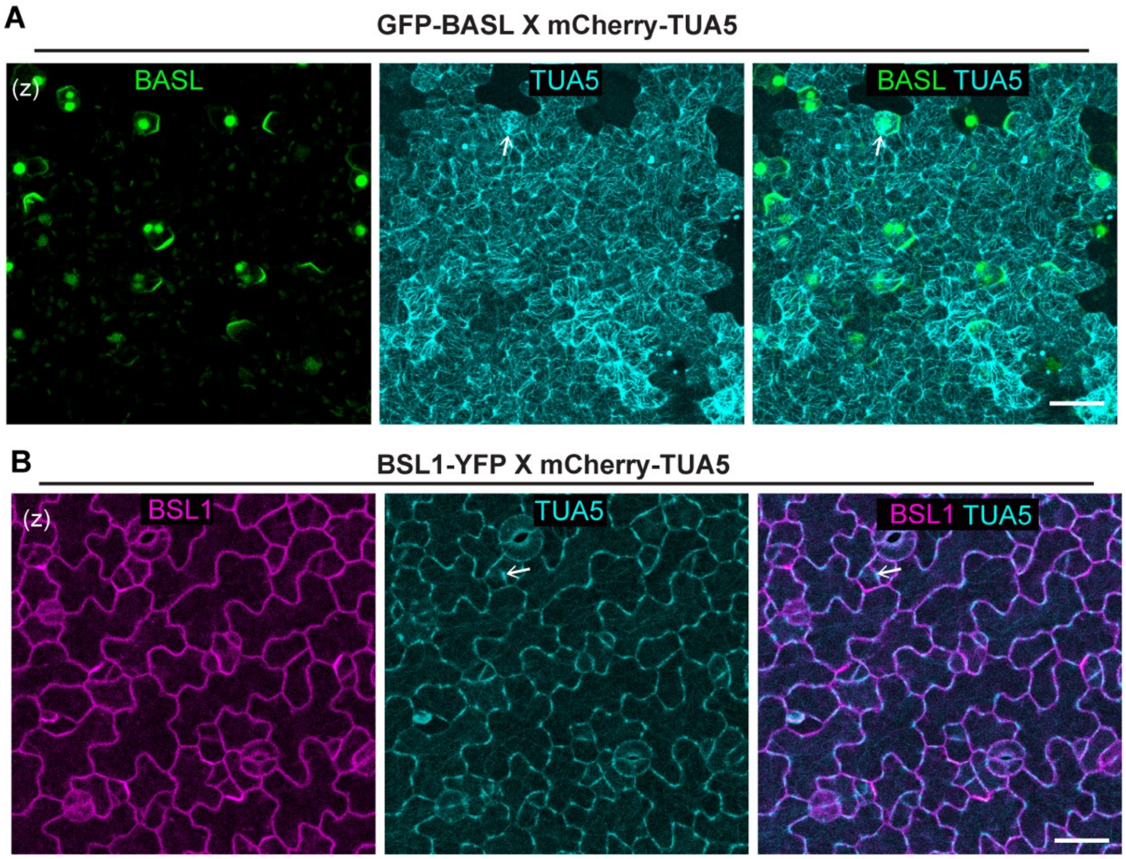


**Figure S3. Determination of Polarization Timing for BASL and BSL1, Related to Figure 3**
(A-B) Co-expression of the native promoter driven GFP-BASL (green)(A) or BSL1-YFP (magenta)(B) with the microtubule marker mCherry-TUA5 (cyan, driven by the ubiquitous *35S* promoter) in 60-hpg adaxial cotyledon epidermis in *Arabidopsis*. The expression of mCherry-TUA5 allows the visualization of the microtubule structure, the preprophase band (PPB) (arrows) that forms at the G2 phase of the cell cycle. Scale bar, 25 μm. (z), images were z-projected.

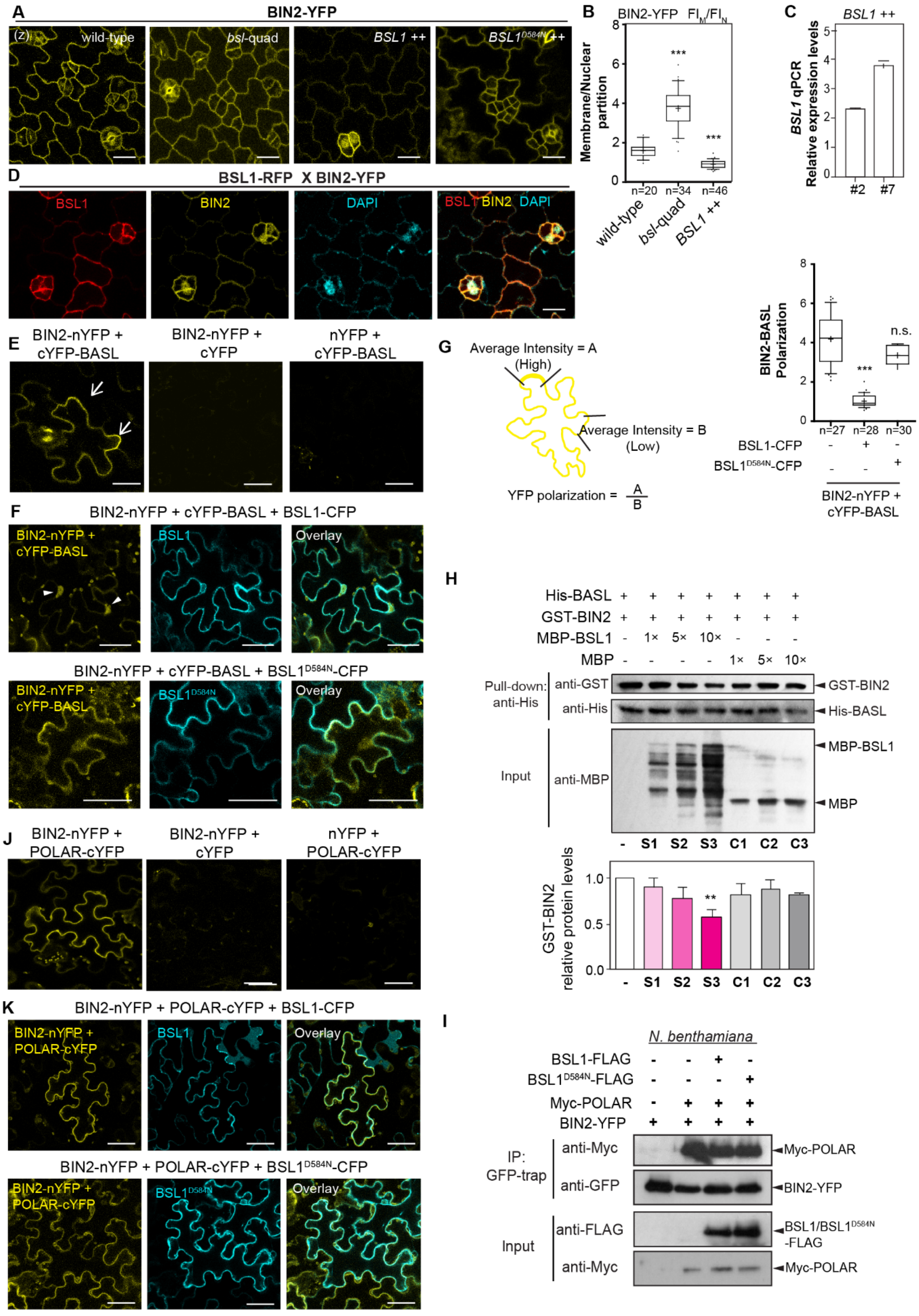

A
BIN2-YFP
(z)
wild-type
bsl-quad
BSL1 ++
BSL1D584N ++
B
BIN2-YFP FIM/FIN
Membrane/Nuclear partition
n=20
n=34
n=46
wild-type
bsl-quad
BSL1 ++
C
BSL1 ++
BSL1 qPCR
Relative expression levels
#2
#7
D
BSL1-RFP X BIN2-YFP
BSL1
BIN2
DAPI
E
BIN2-nYFP + cYFP-BASL
BIN2-nYFP + cYFP
nYFP + cYFP-BASL
G
Average Intensity = A
(High)
Average Intensity = B
(Low)
YFP polarization = A/B
BIN2-BASL Polarization
n=27
n=28
n=30
n.s.
BSL1-CFP
BSL1D584N-CFP
BIN2-nYFP + cYFP-BASL
F
BIN2-nYFP + cYFP-BASL + BSL1-CFP
Overlay
BIN2-nYFP + cYFP-BASL + BSL1D584N-CFP
H
His-BASL
GST-BIN2
MBP-BSL1
MBP
Pull-down: anti-His
anti-GST
anti-His
GST-BIN2
His-BASL
Input
anti-MBP
MBP-BSL1
MBP
S1
S2
S3
C1
C2
C3
GST-BIN2 relative protein levels
J
BIN2-nYFP + POLAR-cYFP
BIN2-nYFP + cYFP
nYFP + POLAR-cYFP
K
BIN2-nYFP + POLAR-cYFP + BSL1-CFP
BIN2-nYFP + POLAR-cYFP + BSL1D584N-CFP
I
N. benthamiana
BSL1-FLAG
BSL1D584N-FLAG
Myc-POLAR
BIN2-YFP
IP: GFP-trap
anti-Myc
anti-GFP
Input
anti-FLAG
BSL1/BSL1D584N -FLAG

**Figure S4. Association of BSL1 with the Polarity Complex Modulates BIN2 Partitioning to the Nucleus, Related to Figure 4**
(A) Representative confocal images show the expression patterns of BIN2-YFP (yellow) in the backgrounds of wild-type *Arabidopsis*, *bsl*-quad mutants, plants overexpressing BSL1 or phosphatase-dead BSL1$^{D584N}$ (both driven by the stomata lineage *TMM* promoter). Scale bars, 20 μm.
(B) Quantification of membrane/nuclear (M/N) partition of BIN2–YFP by measuring fluorescence intensity of z-stacked images. n, number of stomatal lineage cells used for quantification. *** P < 0.0001. One-way ANOVA with Tukey's post hoc test were performed to compare with the values in the wild-type. ***P < 0.0001.
(C) qPCR data show relative expression levels of *BSL1* in two independent overexpression lines (*TMM* promoter) used in this study. Total RNAs were extracted from 3-day-old seedlings. Gene expression levels were normalized by *ACTIN2* and relative expression levels of *BSL1* were compared with the values in the wild-type. Experiments were independently repeated three times. Primers used were listed in Supplemental Table 1.
(D) Co-expression of the *TMM* promoter driven BSL1-RFP (red) with the endogenous promoter driven BIN2-YFP (yellow). The nuclear enrichment of BIN2 was confirmed by DAPI staining (cyan). Scale bar, 10 μm.
(E) BiFC assays show interactions between BIN2 and BASL (yellow) in *N. benthamiana* leaves. Half YFPs (nYFP and cYFP) were used as negative controls. Arrows indicate protein polarization in epidermal pavement cells.
(F) BiFC interaction tests for BIN2-BASL (yellow) in the presence of BSL1-CFP (cyan, top) or BSL1$^{D584N}$-CFP (cyan, bottom). Note, in the presence of BSL1, YFP signals were found diminished along the cell membrane and enriched in the nucleus (arrowheads), suggesting BIN2-BASL interactions were disrupted by the expression of wild-type BSL1 but not by the phosphatase-dead BSL1$^{D584N}$. Representative individual cells in (E-F) were selected from at least three independent experiments. Scale bars in (E-F), 50 μm.
(G) Left: Graphic describes the method for quantification of BASL-BIN2 polarization in the BiFC assays. Confocal images were captured by same settings and z-stacked images were used for measurement of absolute fluorescence intensity. Same lengths were selected from A or B regions and average intensity values were taken to obtain for quantification of protein polarization values. Right: Box plots show quantification of BIN2-BASL polarization in the presence of BSL1-CFP or BSL1$^{D584N}$-CFP. *n*, number of cells. One-way ANOVA followed by Tukey's post hoc test were used to compare with the control. ***P < 0.0001. n.s., not significant.
(H) *In vitro* pull-down assays using recombinant proteins to test the BIN2-BASL interaction in the presence of an increasing amount of MBP-BSL1. His-BASL was used as bait and the amount of GST-BIN2 being pulled down reflects the interaction strength of BIN2-BASL. MBP was used as negative control. Results represent three biological replicates. Histograms (below) show quantification of relative protein levels of BIN2 in the assay above. Results suggest the addition of MBP-BSL1 reduced the amount of BIN2 that interacted with BASL. Data are mean ± SD. Student's *t* test. **P < 0.005.
(I) Co-IP assays using purified fusion proteins produced by *N. benthamiana* leaves show physical association of BIN2-YFP with Myc-POLAR was not influenced by the presence of FLAG-tagged BSL1 or BSL1$^{D584N}$. BIN2-YFP was used as bait. Immunoprecipitated proteins were detected by anti-Myc. Data represent results of experiments repeated for three times.
(J) BiFC assays show protein-protein interaction between BIN2 and POLAR (yellow). Half YFPs (nYFP or cYFP) were used as negative controls.
(K) The interaction of BIN2-POLAR in the BiFC assays (yellow) was not changed by the expression of BSL1-CFP (cyan, top) or BSL1$^{D584N}$-CFP (cyan, bottom). Representative individual cells were chosen from three independent experiments. Scale bars in (J-K), 50 μm.

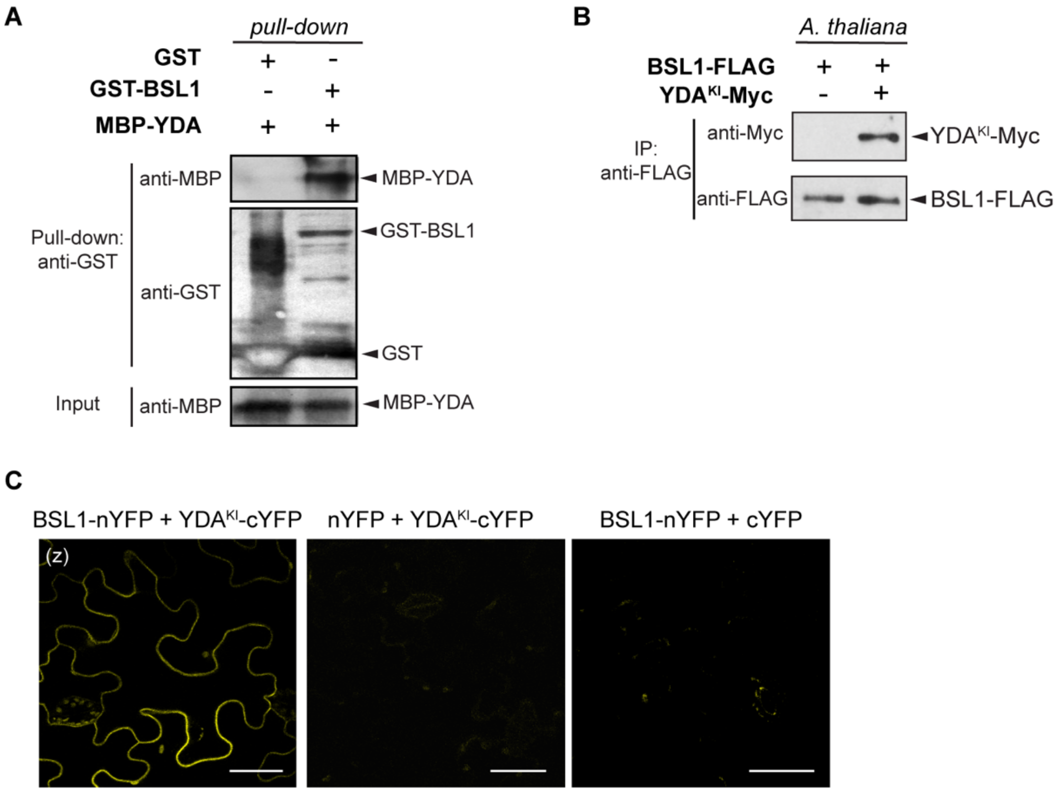


**Figure S5. Association of BSL1 with the Polarity Complex Activates YDA and MAPK Signaling, Related to Figure 5**

(A) *In vitro* pull-down assays using purified recombinant proteins show GST-BSL1 interaction with MBP-YDA. GST (negative control) or GST-BSL1 were used as bait. YDA interaction was detected by anti-MBP.
(B) Co-IP data show *in vivo* interaction between BSL1 and YDA. BSL1-FLAG was used as bait to detect the binding of $YDA^{KI}$-Myc in 5-dpg *Arabidopsis* seedlings.
(C) BiFC assays show the interaction between BSL1 and $YDA^{KI}$ (kinase inactive YDA variant used to suppress active YDA-triggered cell death) occurs at the cell membrane in *N. benthamiana* leaf epidermis. Positive protein-protein interactions were visualized by YFP signal (yellow). Half YFPs (nYFP or cYFP) were uses as negative controls. Scale bars, 50 μm.

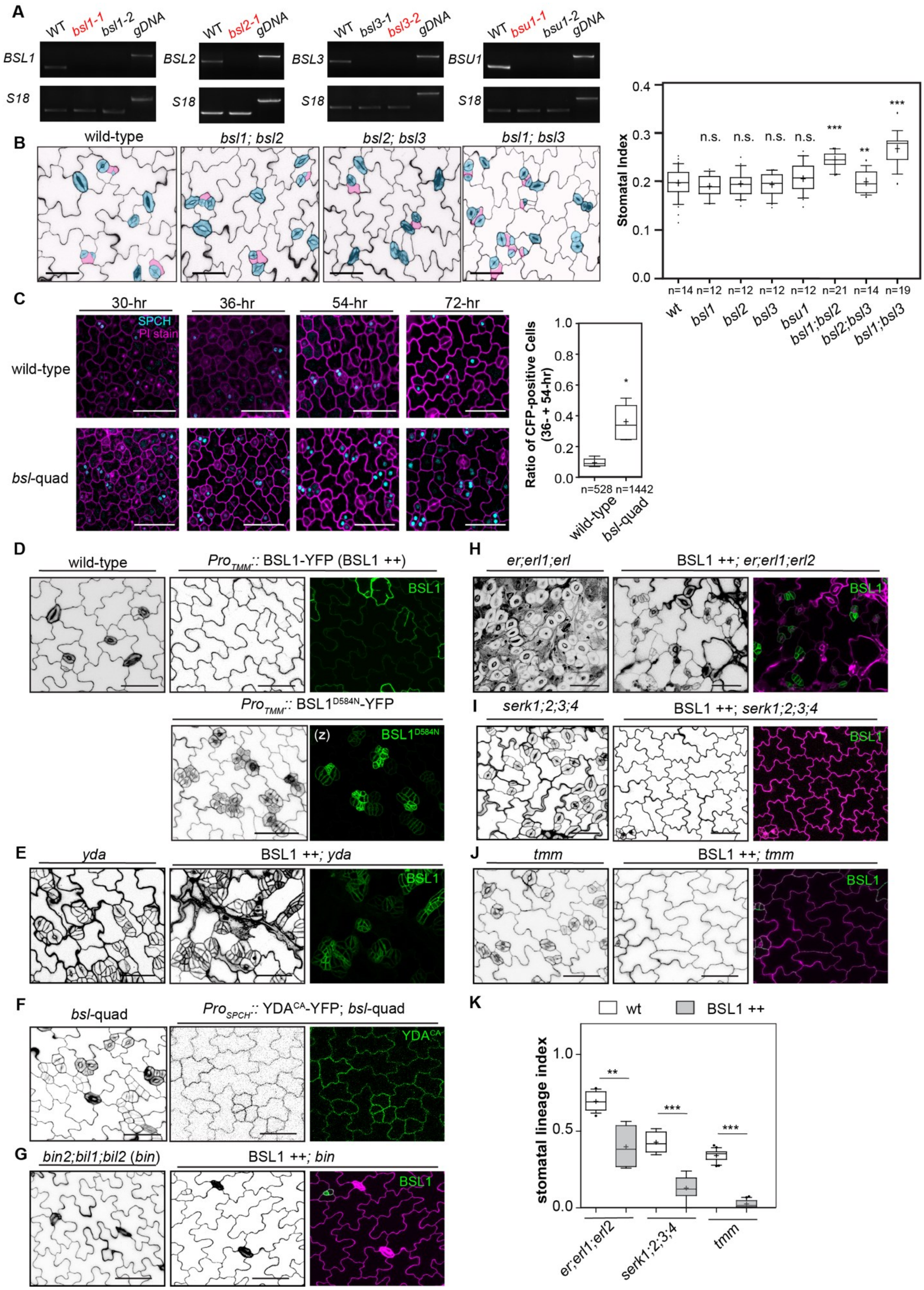

A
WT bsl1-1 bsl1-2 gDNA
BSL1
S18
WT bsl2-1 gDNA
BSL2
S18
WT bsl3-1 bsl3-2 gDNA
BSL3
S18
WT bsu1-1 bsu1-2 gDNA
BSU1
S18
B
wild-type
bsl1; bsl2
bsl2; bsl3
bsl1; bsl3
Stomatal Index
n.s.
***
**
n=14 n=12 n=12 n=12 n=12 n=21 n=14 n=19
wt bsl1 bsl2 bsl3 bsu1 bsl1;bsl2 bsl2;bsl3 bsl1;bsl3
C
30-hr
36-hr
54-hr
72-hr
SPCH
PI stain
wild-type
bsl-quad
Ratio of CFP-positive Cells (36- + 54-hr)
n=528 n=1442
D
wild-type
ProTMM:: BSL1-YFP (BSL1 ++)
BSL1
ProTMM:: BSL1D584N-YFP
(z)
BSL1D584N
E
yda
BSL1 ++; yda
F
bsl-quad
ProSPCH:: YDACA-YFP; bsl-quad
YDACA
G
bin2;bil1;bil2 (bin)
BSL1 ++; bin
H
er;erl1;erl
BSL1 ++; er;erl1;erl2
I
serk1;2;3;4
BSL1 ++; serk1;2;3;4
J
tmm
BSL1 ++; tmm
K
wt
BSL1 ++
stomatal lineage index
er;erl1;erl2
serk1;2;3;4

**Figure S6. BSL Requires BIN2 and YDA to Regulate Stomatal Development in *Arabidopsis*, Related to Figure 6**

(A) Semi-quantitative RT–PCR analysis of single *bsl* T-DNA insertion lines using total RNAs isolated from *Arabidopsis* seedlings at 3-dpg. gDNA, genomic DNA from the wild-type used as template. Primers used for RT–PCR were listed in Supplemental Table 1. Red color indicates the mutant lines used for the generation of high order mutants in this study.
(B) Stomatal phenotype in adaxial cotyledon epidermis of wild-type, *bsl1;bsl2*, *bsl2;bsl3* and *bsl1;bsl3* at 5-dpg. Scale bars, 20 µm. Right, Plot box show Stomatal Index (# stomata relative to # total cells) of the designated genotypes. *n*, number of cotyledons used for quantification of each genetic background. One-way ANOVA with Tukey's post hoc test were used to compare with the wild-type. n.s., not significant. ** $P < 0.005$, *** $P < 0.0001$
(C) Elevated number of cells expressing the stomatal lineage identity marker SPCH in *bsl*-quad mutants. Left: Confocal images (z-projections) show the expression of SPCH-CFP (cyan) in the wild-type (top) and *bsl*-quad mutants (bottom) at different time point (after gemination) during stomatal development. Cell outlines were stained with PI (magenta). Each representative image was selected from 5 comparable cotyledons for each stage. Scale bars, 50 µm. Right: quantification of SPCH-CFP expression in wild-type and in *bsl*-quad mutant. Ratios of CFP-positive cells relative to the total cell numbers in a given areas were calculated. *n*, total number of cells collected from > 20 cotyledons at 36- and 54-hpg. Two-sided *t*-test. * $P = 0.0169$.
(D) Overexpression phenotypes of the *TMM* promoter driven BSL1 (green) or phosphatase-dead BSL1$^{D584N}$ in the stomatal lineage cells in 5-day adaxial cotyledon epidermis. Note the stomatal phenotype generated by BSL1$^{D584N}$ resembled that observed for *bsl*-quad mutants, suggesting a dominant negative effect of BSL1$^{D584N}$.
(E) Overexpression of *BSL1* (green) driven by the *TMM* promoter in the loss of function *yda-3* mutant did not alter the stomatal phenotypes in *yda-3*.
(F) Overexpression of constitutively active YDA (YDA$^{CA}$, green) driven by the *SPCH* promoter in the wild-type (left) and in *bsl*-quad mutants (right). The activities of YDA$^{CA}$ suppressed the loss-of-function phenotypes in *bsl*-quad.
(K) Overexpression of *BSL1* (green) driven by the *TMM* promoter (BSL1 ++) in the loss-of-function mutants *bin2-3;bil1;bil2* (*bin*). *bin* mutations appeared to be epistatic. Cell outlines were stained with Propidium Iodide (magenta).
(H-J) Confocal images show stomatal development (black/white) and protein expression (green) in 5-day adaxial cotyledon epidermis of the designated genotypes. Overexpression of *BSL1* (BSL1 ++, green) were driven by the *TMM* promoter in loss-of-function receptor mutants, i.e. *er;erl1;erl2* (H), *serk1;2;3;4* (I), and *tmm* (J). Cell outlines were highlighted by PI-staining (magenta). Representative images were selected from at least 10 individual cotyledons. Scale bars, 50 µm.
(K) Quantification of Stomatal Lineage Index (SLI) of the genotypes shown in (H-J). More than five cotyledons were scored for each genotype. Statistical analysis was performed with one-way ANOVA followed by Tukey's post hoc test to compare with their respective genetic backgrounds. ** $P < 0.005$, *** $P < 0.0001$.

**Supplemental Table**

**Table S1. List of primers used in this study**

The name and sequence of these primers was displayed in left. The purposes of these primers were listed in right. F, forward; R, reverse.

| Name | Primers | Purpose |
|---|---|---|
| BSL1-F-NotI | GCTCCGCGGCCGCC ATGGGCTCGAAGCCTTG | for pENTR-D1/BSL1-CDS and genomic,overexpression lines, Y2H, BiFc |
| BSL1-R-AscI(-stop) | A GGCGCGCCC GATGTATGCAAGCGAGCTTCTG | |
| BSL2-F-NotI | GCTCCGCGGCCGCC ATGGATGAAGATTCGTCTATGG | for pENTR-D1/BSL2-CDS and genomic,overexpression lines, Y2H, BiFc |
| BSL2-R-AscI(-stop) | A GGCGCGCCC CATCCAAGCCAGAGAACC | |
| BSL3-F-NotI | GCTCCGCGGCCGCC ATGGATTTGGATTCTTCAATG | for pENTR-D1/BSL3-CDS and genomic,overexpression lines, Y2H, BiFc |
| BSL3-R-AscI(-stop) | A GGCGCGCCC TATCCAAGCAAGAGAGC | |
| BSU1-F-NotI | GCTCCGCGGCCGCC ATGGCTCCTGATCAATCTTATC | for pENTR-D1/BSU1-CDS and genomic,overexpression lines, Y2H, BiFc |
| BSU1-R-AscI(-stop) | A GGCGCGCCC TTCACTTGACTCCCCTC | |
| BSL1 promoter-F | GCTCCGCGGCCGCC ACTCAGTTGCATTGAATTTGAC | for pENTR-D1/BSL1promoter-genomic |
| BSL1 promoter-R | GCTCCGCGGCCGCC TGGAAACCACTTTACGGGTATAAATC | |
| BSL2 promoter-F | GCTCCGCGGCCGCC TTATCAAATTGTAGTCCATCCAAG | for pENTR-D1/BSL2promoter-genomic |
| BSL2 promoter-R | GCTCCGCGGCCGCC TATCAAAAAGCTTCAAAAGTGG | |
| BSL3 promoter-F | GCTCCGCGGCCGCC TTCGGTCTTGATGGAACG | for pENTR-D1/BSL3promoter-genomic |
| BSL3 promoter-R | GCTCCGCGGCCGCC ATTTTTCACAACCCTAAATTCGTC | |
| BSU1 promoter-F | GCTCCGCGGCCGCC AAACCACTGACATCTCTTCATC | for pENTR-D1/BSU1promoter-genomic |
| BSU1 promoter-R | GCTCCGCGGCCGCC AACACAATATTTTGTGGTGG | |
| BIN2cds-F-NotI | GCTCCGCGGCCGCC ATGGCTGATGATAAGGAGATG | for overexpression lines, BiFc |
| BIN2pro-F-NotI | GCTCCGCGGCCGCC CTCGGTTATACAATGAGGTTATC | for pENTR-D1/BIN2promoter-genomic |
| BIN2-R-AscI(-stop) | A GGCGCGCCC AGTTCCAGATTGATTCAAGAAG | |
| BSL1D584N-F | CCAAATTGCCCATGGAGATTGCCAAATAC TTTGATAGGA | point mutation |
| BSL1D584N-R | TCCTATCAAAGTATTTGGCAATCTCCATG GGCAATTTGG | |
| BSL1cds-F-SmaI | CC CCCGGG ATGGGCTCGAAGCCTTG | Protein fusion with GST |
| BSL1cds-F-XbaI | GCTCTAGA ATGGGCTCGAAGCCTTG | Protein fusion with MBP |
| BSL1cds-R-SalI(-stop) | GC GTCGAC GATGTATGCAAGCGAGCTTCTG | Protein fusion |

| | | |
|---|---|---|
| BSL2cds-F-EcoRI | CG GAATTC ATGGATGAAGATTCGTCTATGG | Protein fusion with MBP, GST |
| BSL2cds-R-SalI(-stop) | GC GTCGAC CATCCAAGCCAGAGAACC | |
| BSL3cds-F-XbaI | GCTCTAGA ATGGATTTGGATTCTTCAATG | Protein fusion with MBP |
| BSL3-R-SalI(-stop) | GC GTCGAC TATCCAAGCAAGAGAGC | |
| BSU1-F-EcoRI | CG GAATTC ATGGCTCCTGATCAATCTTATC | Protein fusion with MBP, GST |
| BSU1-R-SalI(-stop) | GC GTCGAC TTCACTTGACTCCCCTC | |
| MBP-YDA F | C GAGCTC ATGCCTTGGTGGAGTAAATC | Protein fusion with MBP |
| MBP-YDA R | CC AAGCTT GGGTCCTCTGTTTGTTGATC | |
| pET28a-BASL F | CG GAATTC ATGGCTTCACAGTGGACAATAC | Protein fusion |
| pET28a-BASL(-Stop) R | CC CTCGAG GAATCTACAACATTGGAACC | |
| BIN2-F EcoR1 | CG GAATTC ATGGCTGATGATAAGGAGATG | Protein fusion with GST |
| BIN2cds-R-NotI(-stop) | GCTCCGCGGCCGCC AGTTCCAGATTGATTCAAGAAG | |
| SALK_051383(BSL1) LP | TGATTAAATCTTGTCCACGCC | Mutant genotyping |
| SALK_051383(BSL1) RP | GCTTCATCCGAGAGCTGTATG | |
| SALK_147279(BSL1) LP | GACCTCGAAACTGGAAACCTC | Mutant genotyping |
| SALK_147279(BSL1) RP | TAGGGGTGATTTACCCCAAAC | |
| SALK_055335(BSL2) LP | CATTAGCAAAGTTCTGCCAGC | Mutant genotyping |
| SALK_055335(BSL2) RP | GTTCCAGAGCAGATGGAGATG | |
| WiscDsLox245G08-LP | TGAAGATCGTTGTTGTTGCAG | Mutant genotyping |
| WiscDsLox245G08-RP | AAACTTGTGACATCAGTGGCC | |
| SALK_071689(BSL3) LP | CAAACATTTGAAAGGGTACGATG | Mutant genotyping |
| SALK_071689(BSL3) RP | AAAACATACGAATGCCAGCAC | |
| SALK_072437(BSL3) LP | CCTGCAAAATATCAATGCTTAG | Mutant genotyping |
| SALK_072437(BSL3) RP | TAATGCACTTTTTGGTTTCCG | |
| SALK_030721(BSU1) LP | ACGTTCCACTTCAACATGGAG | Mutant genotyping |
| SALK_030721(BSU1) RP | TCTTTAACCATGCTTCGAACC | |
| SAIL_101_H03 LP | TCAACAAAGGGTCCACAACTC | Mutant genotyping |
| SAIL_101_H03 LP | TGTCCACTTCCTGGTCAAAAC | |
| BSL1 C-LP | GACGACGCTTGGATGCAGGAGCTG | RT-PCR, C to 3'utr 279bp |

| | | |
|---|---|---|
| BSL1 3'utr-RP | CTATACCATTCTCACTCTCTGGT | |
| BSL2-RT LP | GAAGACACATGGATGCAGGAGCT | RT-PCR, C to 3'utr 490bp |
| BSL2-RT RP | CACCTAATCAACCATTACCATTC | |
| BSL3-RT LP | AGAGGATACATGGATGCAGGAGTTA | RT-PCR, C to 3'utr 624bp |
| BSL3-RT RP | CAAACGACCAAACACACCTCTCT | |
| BSU1Ct-RT | CTCCCATCTCATCTTCAG | RT-PCR, C to 3'utr 225bp |
| BSU1 3'utr-R | CCTCTGCCAATACCAAAATAG | |
| BSL1-qPCR-R | CCTCCCTCAATAGCGGTGGCG | qPCR |
